# Influenza A genomic diversity during human infections underscores the strength of genetic drift and the existence of tight transmission bottlenecks

**DOI:** 10.1101/2023.06.18.545481

**Authors:** Michael A. Martin, Nick Berg, Katia Koelle

## Abstract

Influenza infections result in considerable public health and economic impacts each year. One of the contributing factors to the high annual incidence of human influenza is the virus’s ability to evade acquired immunity through continual antigenic evolution. Understanding the evolutionary forces that act within and between hosts is therefore critical to interpreting past trends in influenza virus evolution and in predicting future ones. Several studies have analyzed the longitudinal patterns of influenza A virus genetic diversity in natural human infections to assess the relative contributions of selection and genetic drift on within-host evolution. However, in these natural infections, within-host viral populations harbor very few single nucleotide variants, limiting our resolution in understanding the forces acting on these populations in vivo. Further, low levels of within host viral genetic diversity limit the ability to infer the extent of drift across transmission events. Here, we propose to use influenza virus genomic diversity as an alternative signal to better understand within and between host patterns of viral evolution. Specifically, we focus on the dynamics of defective viral genomes (DVGs) which harbor large internal deletions in one or more of influenza virus’s eight gene segments. Our longitudinal analyses of DVGs show that influenza A virus populations are highly dynamic within hosts, corroborating previous findings based on viral genetic diversity that point towards the importance of genetic drift in driving within-host viral evolution. Further, our analysis of DVG populations across transmission pairs indicate that DVGs rarely appeared to be shared, consistent with previous findings indicating the presence of tight transmission bottlenecks. Our analyses demonstrate that viral genomic diversity can be used to complement analyses based on viral genetic diversity to reveal processes that drive viral evolution within and between hosts.

**Author summary:** During viral replication within infected individuals, different types of mutation can occur. Point mutations introduce genetic diversity, with some sites in the viral genome becoming polymorphic at the viral population level. Structural mutations that result in deletions in the viral genome can also occur, contributing to genomic diversity in the viral population. Defective viral genomes (DVGs) are a particular type of this genomic diversity that are incapable of establishing productive infection on their own, but persist during infection through cellular coinfection with infectious virus. Here, we first identify DVGs in deep sequencing data from natural infections of influenza A virus subtype H3N2 and then use these DVGs to inform our understanding of within-host viral evolution and transmission dynamics. By analyzing DVG populations over time within infected individuals and between transmission pairs, we find that influenza A virus populations are shaped by genetic drift and tight transmission bottlenecks. Our results add to our understanding of influenza genomic diversity within-and between-hosts and demonstrate the possibility of considering the full suite of viral diversity to understand drivers of viral evolution.

## Introduction

Influenza A virus is one of the most common viral pathogens worldwide. Despite relatively widespread vaccination, influenza infection results in over 20,000 deaths and $3.7 billion in direct medical costs each year in the United States alone [1]. One of the contributing factors to the virus’s widespread circulation is its ability to rapidly evolve to evade natural and vaccine-derived immunity [2, 3, 4, 5, 6]. This so-called “antigenic drift” allows the virus to continually replenish its pool of susceptible hosts by reinfecting hosts who already harbor immunity to previously circulating strains. Understanding how new antigenic variants evolve within single hosts and ultimately sweep the population is important for vaccine strain selection [7, 8, 9] and informing the development of vaccines that are more robust to viral evolution [10].

The evolution of antigenically diverse lineages of influenza virus is possible because there is diversity in the viral population on which selection can act. This diversity is generated by errors made by the viral polymerase during replication within single hosts and is impacted by the evolutionary forces that occur within hosts. Population bottlenecks that occur during transmission between hosts further shapes this viral diversity [11]. Analyzing these dynamics can therefore provide insights into the evolution acting within-and between hosts.

Deep sequencing data can be used to characterize patterns of viral genetic diversity within individual hosts. To do so, sequencing reads are aligned to a reference genome and used to estimate the frequency of each nucleotide at each site of the viral genome. By analyzing the frequencies of these intrahost Single Nucleotide Variants (iSNVs) across multiple time points one can determine whether selection is acting on specific mutations or whether genetic drift dominates within host evolution [12]. Finally, by comparing iSNV frequencies in source and recipient hosts in transmission pairs one can estimate how many viral particles seeded infection in the recipient [13].

Previous analyses of this type have revealed that selection during an acute natural influenza A virus infection is relatively weak. Positive selection acting on known antigenic escape mutations is not apparent and the presence of prior immunity has little impact on the amount of observed within host genetic diversity [14, 11]. It has been suggested that this may be due to mismatch in the timing between viral population growth and the immune response [15]. While evidence for positive selection is limited (with the exception of infections in children [16]), purifying selection does appear to act within these infections and contribute to shaping in vivo influenza A virus populations [11].

Our understanding of influenza A virus evolution within individuals and between transmission pairs stems from comprehensive analyses of many different individuals and transmission pairs. This is because within any given acute infection, there is limited viral genetic diversity. At a 2-3% variant calling threshold, the number of iSNVs identified in a viral sample is generally fewer than 15, and most of these iSNVs occur at low frequency [14, 11]. This limited genetic diversity hinders our ability to robustly characterize the evolutionary forces acting on these populations and results in considerable uncertainty in our inferred contributions of selection and drift to within and between host evolution. Here, we propose using an alternative signal to study the evolutionary dynamics of viral populations within and between hosts. Specifically, we propose focusing on viral genomic diversity that is generated during infection, in the form of influenza defective viral genomes (DVGs).

DVGs (here used synonymously with deletion-containing viral genomes, DelVGs [17]) harbor a large internal deletion in at least one of the eight segments of the influenza A virus genome. As a result, virions with a DVG are incapable of replicating on their own. However, through coinfection of a cell with an infectious “wild-type” virus, they can proliferate throughout an infection [17]. The process by which coinfection rescues cellular infection with DVGs is similar to the process by which coinfection can rescue infection by virions with incomplete genomes [18, 19] except that instead of missing entire segments, DVGs harbor truncated copies of gene segments. Individual DVG segments can be identified by the genomic sites at which these deletions occur (“DVG species”). Due to the reliance on coinfection, we expect the evolutionary forces acting on DVG populations to mirror those acting on the wild-type viral population. For example, if positive selection is acting on a specific viral mutation, then DVG species that coinfect with wild-type viruses harboring this mutation will appear to have an evolutionary advantage over the DVG species which coinfect with cells lacking this beneficial mutation.

This is analogous to the process of genetic hitchhiking, in which loci that are linked to beneficial mutations will increase in frequency [20]. Hitchhiking can occur due to physical linkage on a gene segment or through spatial structure that maintains linkage disequilibrium. Given the extent of spatial structure in within-host influenza virus infections [21], we therefore expect linkage to be strong not only within gene segments but also between them, consistent with the limited effective reassortment between gene segments that has been found in a longitudinally-sampled human influenza challenge study [22]. With linkage between DVGs and co-circulating wild-type viruses, the dynamics of DVGs may thus offer a complimentary signal of evolutionary processes occurring within hosts to that which is presented by viral genetic diversity in the form of iSNVs.

It has been proposed that DVGs may also be transmitted between hosts [23]. Transmission of DVGs between hosts must rely on coinfection of recipient host cells by both wild-type and DVG virions. *A priori*, this is expected to be unlikely under the assumption that wild-type and DVG virions infect host cells at random during transmission, given the large number of susceptible host cells. However, it is thought that some viruses, influenza included [24], may form so-called collective infectious units [25]. In these collective units, virions aggregate together, presumably increasing the probability of cellular coinfection. This may play a role in how influenza viruses are able to transmit at all, given the high proportion of virions harboring incomplete viral genomes [18, 19]. If viral aggregates form, and wild-type virions within these aggregates are genetically identical or near identical due to the spatial structuring of the source individual’s viral population, then DVGs may provide more resolution into the characterization of transmission bottlenecks.

Here, we apply a recently developed bioinformatic pipeline [26] to previously published deep sequencing data from 217 clinical samples of influenza A H3N2 infections from 168 naturally infected, otherwise healthy, individuals from a cohort study [11]. Longitudinal samples are available for 48 of these 168 individuals, allowing us to examine patterns of DVG diversity within hosts over their course of infection. Deep sequenced virus samples are available for 39 epidemiological transmission pairs, allowing us to use DVG diversity to characterize the transmission bottleneck between sources and recipients.

## Materials and methods

### Data source

All clinical and sequencing data were previously published as part of [11] and all epidemiological and laboratory methods are described in detail in the original publication. In short, the HIVE cohort at the University of Michigan School of Public Health queries participating households weekly during the months of October through May for symptoms of respiratory illness. Individuals with symptoms were sampled via a combined nasal and throat swab by the research team. During the 2014-2015 season individuals were also instructed to take a self-or parent-collected nasal swab at symptom onset.

cDNA was amplified from samples testing positive for influenza virus using the SuperScript III ONe-Step RT-PCR Platinum Taq HiFi Kit and the universal influenza A primers [27]. Sequencing libraries were prepared from 300-400bp sheared cDNA fragments and barcoded libraries were further purified by isolation of a 300-500 bp band using gel isolation. 2 × 125 nucleotide paired end reads were generated on an Illumina HiSeq 2500. Samples with input titers between 10^3^ and 10^5^ genomes/*µl* were sequenced in replicate. For each sequencing run PCR amplicons derived from eight clonal plasmids of the circulating strain were sequenced on the same HiSeq flow cell as the clinical samples.

Influenza A H3N2 (as identified by a library labelled “perth,” “hk,” or “vic”) sequencing reads were downloaded from the National Library of Medicine (NLM) National Center for Biotechnology Information (NCBI) Sequencing Read Archive (SRA) BioProject PRJNA412631 [11] using the fasterq-dump utility available as part of the SRA Toolkit (https://github.com/ncbi/sra-tools).

### Transmission analyses

We used the same household and unlinked pairings as reported in [11]. Pairs of epidemiologically unlinked samples (“unlinked pairs”) were generated by randomly assigning pairs of samples within each season, excluding samples from the same household. Epidemiologically linked pairs (“transmission pairs”) were identified as in [11] from pairs of individuals from the same household who were infected with influenza viruses more similar than 95% of unlinked pairs, as measured by the L1-norm. When multiple samples were available from the same host, we used the set of DVGs present in all samples for a given host, taking the maximum relative read support observed across samples.

### DVG identification

DVGs were identified using a modified version of the pipeline presented in [26]. Reference genomes were used in accordance with those used in [11]: GenBank CY121496-503 was used for samples collected as part of the 2010-2011 or 2011-20112 season, GenBank KJ942680-8 was used for samples collected during the 2012-2013 season, and GenBank CY207731-8 used for samples collected during the 2014-2015 season. To aid in the identification of DVGs with breakpoints near the 5’ or 3’ end of a segment we added a 210nt poly-A pad to the 5’ and 3’ end of each segment. The analysis pipeline was run in Nextflow v19.01.0 [28].

For quality control sequencing reads were first trimmed using Trimmomatic v0.39 [29] in Phred33 mode using the TruSeq3-PE-2 adapters allowing for 2 seed mismatches, a palindromeClipThreshold of 15, and a simpleClipThreshold of 10, scanning the read with a 3 base sliding window and cutting when the average quality falls below 20, removing leading and trailing bases with quality less than 28, and removing reads less than 75 nucleotides in length. Next Kraken2 v.2.0.7-beta [30] with the k2_pluspf_16gb database in paired end mode was used to categorize each read. The extract_kraken_reads.py script from Kraken Tools (https://github.com/jenniferlu717/KrakenTools) was used to filter only for reads assigned to influenza A virus (taxonomic id 11320) or any children taxa. Quality control on the filtered reads was conducted with FastQc v0.11.9 [31].

The paired end reads were concatenated into a single fastq file and aligned using Bowtie2 v.2.4.2 [32] in end-to-end mode with a minimum scoring scheme of L,0,-0.3. End-to-end mode disallows soft-clipping of reads which is needed to align reads that span DVG junction sites. Therefore, reads which do align in end-to-end mode include all wild-type viral reads as well as reads from the 5’ and 3’ ends of DVGs which do not span the deletion junction site (hereby referred to as non-DVG supporting reads). The number of reads that aligned to each segment in the step was calculated using Samtools v1.13 with htslib v.1.13 [33] by first sorting by name, then adding mate score tags with fixmate, sorting again by coordinates, marking and removing duplicates with markdup and finally tabulating reads using idxstats.

Sequencing reads which did not align in end-to-end mode include the DVG deletion spanning reads and were thus used as input to ViReMa v0.25 [34] with a seed length of 25 nucleotides, tolerating 1 mismatch in the seed alignment, and allowing up to 8 mismatches at the 5’ and 3’ end of an alignment. To exclude small indels we removed any DVG-supporting reads which support a deletion of less than 20 nucleotides. Duplicate DVG supporting reads were also removed with ViReMa. Bowtie v1.0.0 [35] was used within ViReMa. Due to the size filter in the sequencing protocol described above, we are unable to identify any DVGs that are less than 300 nucleotides in length.

As discussed in [26], identifying the precise DVG breakpoint from sequencing data can be impossible when there are nucleotide repeats on either side of the deletion junction, as is prone to occur [17]. ViReMa includes a “DeFuzz” feature which allows users to force the reported junction site to either the 5’ or 3’ end of a given read. However, because DVGs may be supported by reads either in the forward or reverse direction, this behavior results in disparate reporting for the same DVG depending on the supporting read direction. Therefore, we slightly modified ViReMa’s underlying AddToDict function to force the reported junction sites to the 5’ or 3’ end of the reference genome, instead of the supporting read. All DVG junctions reported here have been DeFuzz’d to the 3’ end of the reference genome. ViReMa results were parsed in Perl v5.26.2 using the summary scripts included as part [26]. Specific DVG segments are identified by the genomic location (1-indexed) of the nucleotides flanking the deleted nucleotides. For example, PB2 100_800 represents a DVG generated from the PB2 segement in which nucleotide 101 through 799 have been deleted.

### DVG quality filtering

DVGs for each clinical sample as well as the plasmid control for each sequencing run were further parsed using Python v3.10.8 [36] with Pandas v1.5.2 [37] and Numpy v1.23.5 [38]. The reported junction breakpoints from the three sets of reference genomes were reconciled by pairwise aligning the segments from each reference using MAFFT v7.6.4 [39] and creating a mapping between the coordinates for each individual reference and a universal coordinate system. DVGs breakpoints are reported in the universal coordinate system. For each identified DVG, we calculated its relative read support by normalizing the number of DVG supporting reads by the sum of the DVG supporting reads and the non-DVG supporting reads for that segment.

Notably, this relative measure is imperfect because reads from the 5’ and 3’ portions of the genome which are conserved in DVGs cannot be reliably assigned to having been derived from a DVG or wild type viral genome. Furthermore, because longer segments harbor more internal coding nucleotides that can be deleted by DVGs without affecting packaging, this metric may bias the relative support of DVGs on longer segments downwards. Nevertheless, this measure should be a reasonable means of adjusting for variable sequencing depth, particularly when comparing segments of similar length (such as the three polymerase segments).

For each sequencing run, we first excluded any DVG species (as identified by their junction breakpoints) observed in their respective plasmid control. Where technical sequencing replicates from the same biological sample were available we included only DVGs present in both replicates. In these cases, the mean of the relative support values from each replicate was used in downstream analyses. Based on the read support of DVGs present in the plasmid controls, we included only DVGs supported by at least 10 sequencing reads in downstream analyses.

### Statistical analyses and visualization

All statistical analyses were done in Python. Statistical tests were conducted using Scipy v1.9.3 [40] and regression modeling was done using statsmodels v0.14.0 [41]. All visualization was done in Python using Matplotlib v3.6.2.

### Data and code availability

All raw sequencing data is available from the Center for Biotechnology Information (NCBI) Sequencing Read Archive (SRA) BioProject PRJNA412631 [11]. Additional metadata is available as part of [11] as well on their repository at https://github.com/elifesciences-publications/Host_level_IAV_evolution. All additional analysis and visualization code is available at https://github.com/m-a-martin/influenza_dvg.

## Results

### Limited DVGs in plasmid controls

To determine the number of spurious DVGs introduced by the sequencing protocol, we first queried the reads generated from the plasmid controls for each sequencing run for DVGs. This analysis acts as a negative control as these DVG reads are generated from clonal plasmids containing full-length viral genome segments and thus should not contain any DVGs. While we observed many (median [sd]: 44.00 [49.54]) unique DVGs in each plasmid control, encouragingly, these DVGs were almost exclusively present at very low read support (Fig 1), 1.00 [0.52]). The vast majority (*>*99.9%) of DVGs identified in plasmid controls are supported by *<*10 reads. Consequently, we required DVGs to be supported by a minimum of 10 sequencing reads for all downstream analyses to remove spurious DVGs introduced by the sequencing protocol.

**Fig 1.**
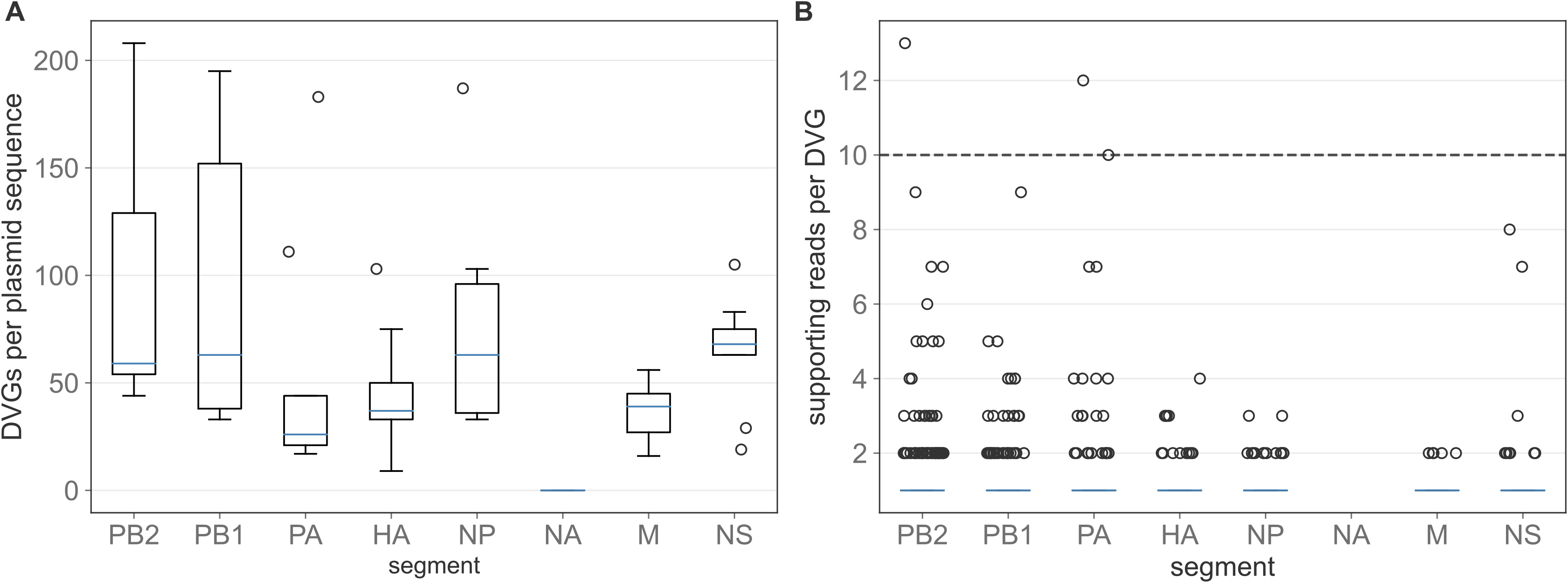
DVGs identified in the plasmid controls. Boxes extend to the limits of the inter-quartile range (IQR), and whiskers extend to 1.5 IQR below and above the 1st and 3rd quartile, respectively. Outliers are shown as dots beyond the range of the whiskers. Blue line represents median values. (A) Number of unique DVGs identified per sequenced plasmid per genome segment. (B) Number of supporting reads per DVG per sequenced plasmid per genome segment. Dotted horizontal line at *N* = 10 reads represent threshold used for downstream analyses.

### DVGs are observed readily in clinical samples

As opposed to the plasmid controls described above, DVGs are observed readily in the clinical samples (Fig 2A). We observe at least one DVG above the supporting read threshold of 10 in all 217 of the clinical samples. DVGs are observed most abundantly on the PB2, PB1, PA, and NS segments (Fig 2A,B, Mann-Whitney U test comparing the relative read support of PB2, PB1, PA, and NS DVGs v. HA, NP, NA, and M DVGs *p*-value *<* 1e*-*4). However, when looking at the number of unique DVGs as defined by their junction breakpoints (Fig 2C), we see a slightly different pattern: PB2, PB1, and PA harbor many more unique DVG species than other segments. NS contains very few unique DVGs (Mann-Whitney U test comparing the number of DVGs per sample observed on the PB2, PB1, PA segments v. on the HA, NP, NA, M, and NS segments *p*-value*<* 1e*-*4). This pattern is consistent with the presence of one or very few unique NS DVGs which are abundantly supported. Further investigation revealed that a single NS DVG (316_545) is identified in 197 out of 217 clinical samples whereas the next most prominent DVG is identified in only 31 samples. Out of 7,481 unique DVGs, 6,098 are observed in only a single sample (S1 Fig). Our finding that DVGs primarily proliferate on the polymerase segment is consistent with previous *in vitro* [17, 42] and *in vivo* [23] analyses. Based on our finding that (with the exception of NS), most observed DVGs fall on the polymerase segments, we here limit our downstream analyses to DVGs found on PB2, PB1, and PA.

**Fig 2.**
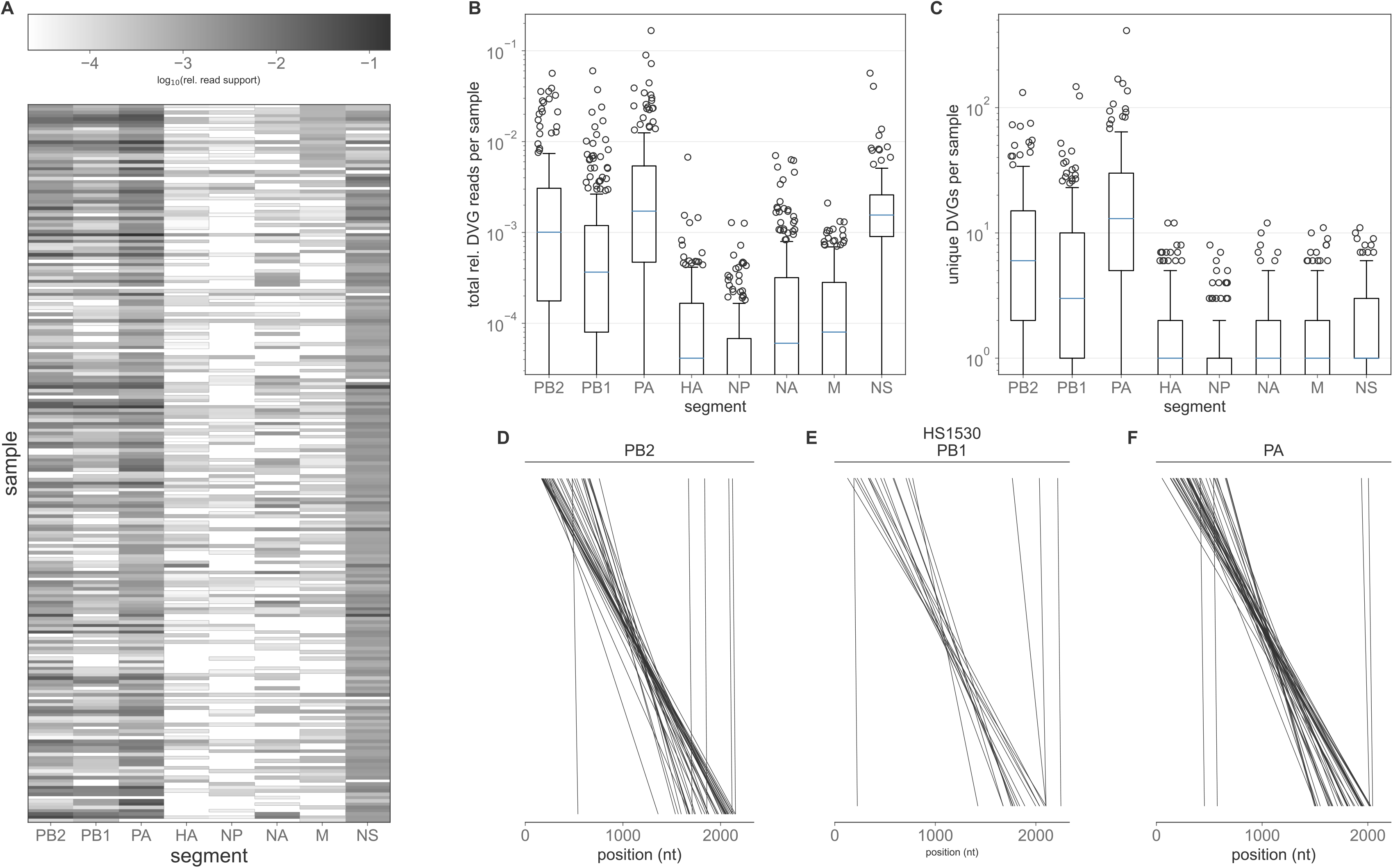
DVGs identified in clinical samples. (A) Heatmap showing the prevalence of DVGs in each sample (rows), by gene segment (columns). DVG prevalence is quantified by the total number of reads that support DVGs, relative to the total number of reads. (B) Total relative DVG reads per sample per segment. (C) Number of unique DVGs per sample per segment. In (B) and (C), blue lines in the boxplots show the median value for each segment, box extends to the limits of the IQR, and whiskers extend to 1.5 IQR below and above the 1st and 3rd quartile, respectively. Outliers are shown as dots beyond the range of the whiskers. Y-axis is truncated as lower quantiles extend to 0. (D,E,F) Breakpoints of all PB2 (D), PB1 (E), and PA (F) DVGs identified in representative sample HS1530. Each line connects the last undeleted base on the 5’ end of the DVG and the first undeleted base on the 3’ end of the DVG.

Canonically, the generation of influenza DVGs results in the deletion of the internal coding region for each segment and the conservation of the 5’ and 3’ termini, which are thought to be needed for virion packaging [43]. To assess whether our identified DVGs followed this pattern we mapped the deletion junction sites onto the reference genome (Fig 2D,E,F, S2 Fig). This analysis reveals that the majority of observed DVGs do indeed result in the deletion of the internal portion of each segment, providing confidence that our analysis pipeline is accurately identifying defective genomes, that is, those which are incapable of establishing productive infection in the absence of wild-type virus. Our analysis also identified a small number of polymerase DVGs harboring relatively small deletions. A comprehensive analysis of the number of deleted nucleotides in each of the observed polymerase DVGs reveals a highly bi-modal pattern (S3 Fig). To ensure that our analyses were based solely on truly defective genomes and not those harboring small indels with minimal fitness effects, we implemented an additional empirical filtering step to remove any DVGs with fewer than 500 deleted nucleotides.

### DVG populations are dynamic

Our primary goal in this study was to assess how polymerase DVG populations change over time during the course of single infections and during transmission between hosts. To assess the former of these two goals, we first analyzed both the total relative polymerase DVG read support as well as the number of unique polymerase DVGs on a per sample basis as a function of the days post symptom onset, which is used in the absence of data on time since infection (Fig 3A,B). Our results indicate that both the total relative DVG reads and the number of unique DVG reads per sample do not vary based on the number of days post symptom onset (Kruskal-Wallis H-test *p*-value=0.08 and 0.06, respectively). The lack of temporal pattern in the number of unique DVG reads is consistent with the lack of temporal pattern in the number of iSNVs identified in these same samples [11].

**Fig 3.**
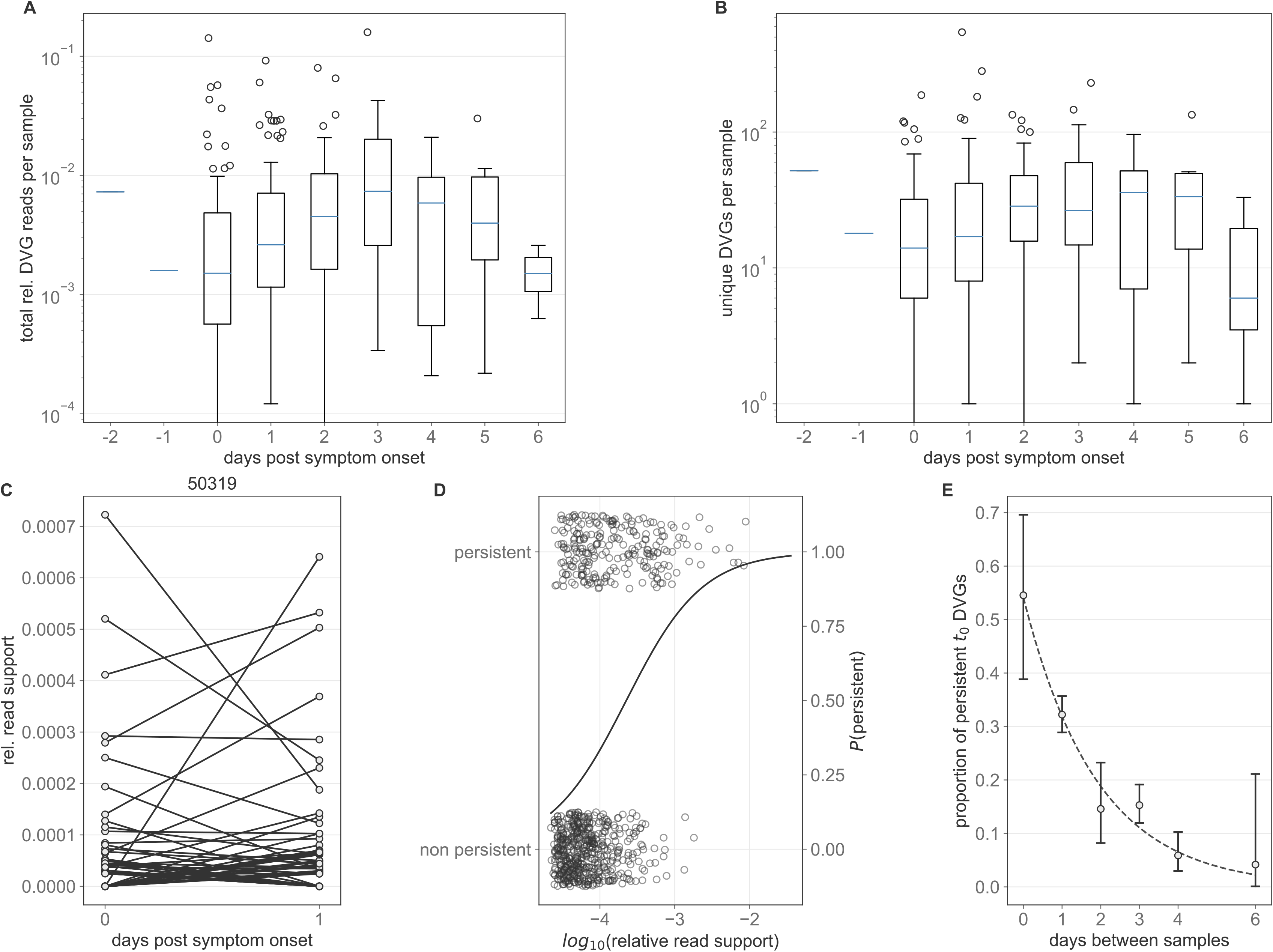
**Longitudinal DVG dynamics.** Polymerase DVG dynamics over the course of infection. (A) Total relative polymerase DVG reads per sample as a function of the number of days post symptom onset that a given sample was taken. (B) Number of unique polymerase DVGs per sample as a function of the number of days post symptom onset that a given sample was taken. (C) Longitudinal DVG dynamics in from representative individual 50319. Lines connect the relative read support of a given polymerase DVG in each of the two samples taken. (D) Relative read support of *t*_0_ DVGs which do and do not persist in the *t*_1_ for longitudinal samples sampled from the same subject 1 day apart. Solid line represents the estimated probability of a DVG with a given read support persisting from a logistic regression. (E) Proportion of all DVGs observed in a given *t*_0_ sample which is also observed in the corresponding *t*_1_ sample as a function of the number of days between when those samples were taken. Whiskers extend to the exact binomial confidence intervals for a given proportion. In all boxplots (A, B), blue line represents the median value for each segment, box extends to the limits of the IQR, and whiskers extend to 1.5 IQR below and above the 1st and 3rd quartile, respectively. Outliers are shown as dots beyond the range of the whiskers.

This lack of temporal pattern indicates that either DVG populations are generated prior to the earliest samples in this dataset and then persist throughout infection or that there is continual turnover (generation and loss) of novel DVG species throughout infection. To determine which of these two possibilities explain the lack of temporal pattern in DVGs, we compared the DVG populations present at multiple time points, subject to availability. In general, DVG populations are highly dynamic over time (Fig 3C, S4 Fig) and many DVGs do not persist across time points. This implies that there is continuous *de novo* DVG generation and extinction of previously generated DVGs during infections.

We first hypothesized that the probability that a DVG persists across longitudinal time points is dependent on the frequency of that DVG at the first time point. Using logistic regression, we tested whether the log-odds that a DVG identified in the first sample of a pair was still present in the second sample of the pair depended on the log_10_ relative read support of the DVG identified in the first sample. Among samples taken 1 day apart (N=24), we find that read support is significantly associated with the log odds of DVG persistence (*p*-value*<* 1e *-* 4, Fig 3D, S1 Table) such that a 1 log-unit increase in the relative read support is associated with an odds ratio of persistence of approximately 7. Specifically, between a log10 relative read support of −4 and −2 the probability of DVG persistence increases from 0.33 to 0.96.

Our longitudinal data also include samples that were taken at a range of time intervals, from 0 to 6 days between samples. These longitudinal data include self-or parent-collected nasal swabs at illness onset and combined nasal and throat swabs at a later visit to the research clinic (see [11]). Thus, we are able to correctly specify the temporal ordering of samples even among those taken on the same day. To evaluate the impact of the time between samples on the probability of DVG persistence, we examined how quickly DVGs that were identified in one sample were no longer observed in a later sample. To this end, we identified the unique DVG species present in the earliest sample (*t*_0_) from each individual with two samples in this dataset and calculated the proportion of these which were present in their later (*t*_1_) sample. By plotting this proportion as a function of the number of days between the *t*_0_ and *t*_1_ samples, we found that this proportion significantly depends on the number of days which have elapsed between *t*_0_ and *t*_1_ (*x*^2^ test of independence p-value*<* 1e *-* 4, Fig 3E). Specifically, the further apart in time that the two samples from a host were taken, the fewer *t*_0_ DVGs persisted in the *t*_1_ sample. Only approximately 33% of DVGs present at *t*_0_ were present a day later and only 4% of *t*_0_ DVGs were present in samples taken six days later. These result are further supported by a multivariate logistic regression model incorporating both relative read support and time between samples as a categorical variable (S5 Fig, S2 Table). In this model both relative read support and time between samples are significant predictors of the probability of DVG persistence (all *p*-values≤ 1e *-* 3).

Taken together, our results indicate that within-host DVG persistence is governed primarily by the time between samples and the DVG quantity within a sample. These findings are consistent with a model in which within-host viral populations are dominated by genetic drift as opposed to selection acting on specific subsets of the viral population.

### DVGs support the existence of a tight transmission bottleneck

Viral evolution is shaped not just by the dynamics which operate within hosts but also by those which operate between hosts, including at the stage of transmission. Previous analyses based on viral genetic variation have revealed that the transmission bottleneck of influenza A is quite small, on the order of one to two virions [11]. This results in a significant loss of genetic diversity during transmission such that nearly all genetic variation observed within a host is likely to have been generated *de novo* following transmission.

To determine whether genomic diversity supports these conclusions, we wished to evaluate how DVG populations compare between known source and recipient pairs. To do this, for each identified transmission pair we tabulated the number of unique DVGs identified in the source that were also identified in the recipient (and thus could have been transmitted from source to recipient). We compared this to a null distribution of epidemiologically unlinked samples to account for the *de novo* generation of identical DVGs within each host.

We found that epidemiologically unlinked samples generally share very few polymerase DVGs, on the order of zero to one (mean [sd] = 0.39 [1.03], Fig 4A). This indicates that *de novo* DVG generation on polymerase segments does not result in the repeated emergence of a small number of specific DVGs. Amongst transmission pairs, we similarly found that generally very few DVGs are shared between individuals (0.54 [1.23]). The number of DVGs shared between random pairs and household pairs was found to not be be significantly different (Mann-Whitney U test *p*-value=0.788), indicating that transmission pairs are not likely to share more DVGs than unlinked pairs. This result indicates that DVG species that are shared between individuals in a transmission pair likely arose de novo within the donor and within the recipient.

**Fig 4.**
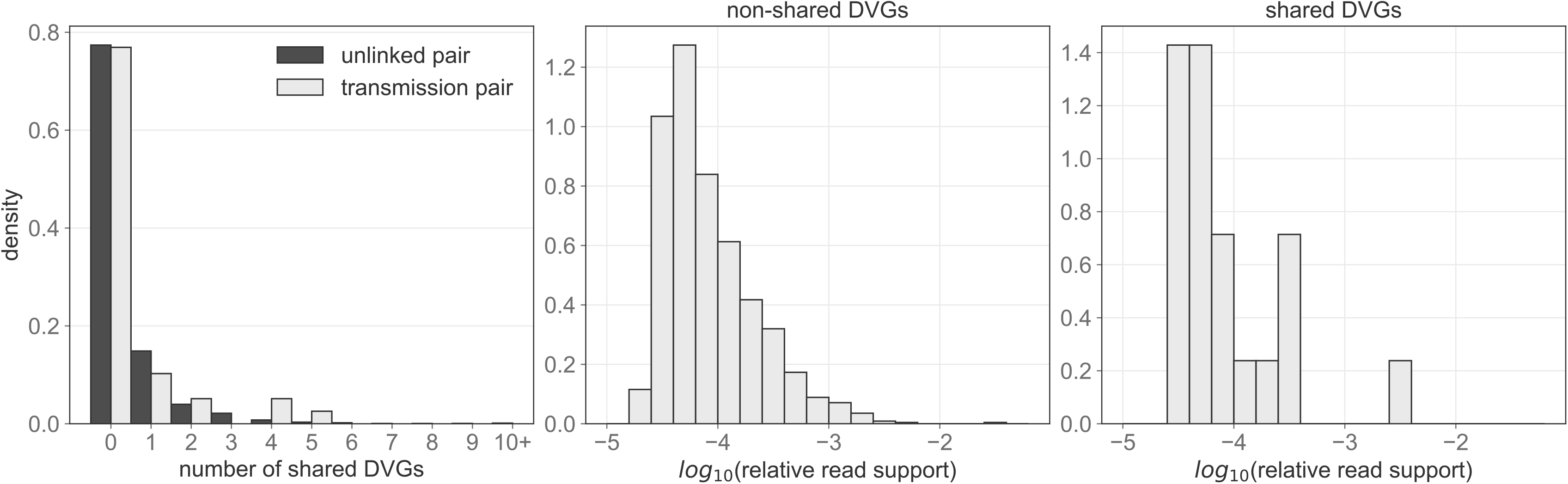
Polymerase DVGs shared between pairs of individuals. (A) Histogram showing the number of DVGs shared between epidemiologically unlinked pairs (dark grey) and between transmission pairs (light grey). XI- axis has been truncated to 10 for visualization purposes, despite a long tail on the unlinked pair distribution. (B) Relative read support in the source individual of DVGs that are not shared between donor and recipient individuals. (C) Relative read support in the source individual of DVGs that are shared between donor and recipient individuals.

While transmission pairs do not tend to share more DVGs as compared to random pairs, we do observe 21 shared DVGs from 9 transmission pairs. If these DVGs were shared due to transmission as opposed to *de novo* generation we would expect them to be disproportionately present at higher relative read support in the donor host relative to DVGs which were not transmitted. However, we observe that the relative read support of shared DVGs (2.2e*-*4 [(5.4e*-*4]) is comparable to that of non-shared DVGs (1.9e*-*4 [(1.1e*-*3]) We found no significant difference between the relative read support in the donor host of shared and non-shared DVGs (Fig 4B,C, Mann-Whitney U *p*-value=0.648). Taken together, these results support the conclusion from iSNV analyses that transmission bottleneck sizes are very small; DVG species do not appear to be transmitted from a donor to a recipient, a pattern that is consistent with a small bottleneck that only results in the establishment of a small number of infectious viral particles in a recipient host.

## Discussion

Understanding how influenza viral populations evolve within and between hosts is key to understanding how viral evolution proceeds on the host population scale. This is relevant, for example, in learning how new antigenic variants arise and ultimately sweep the host population. Previous work has used genetic diversity, or diversity in the form of single nucleotide variants, to better understand the evolutionary forces acting on virus populations within hosts. These analyses have found that positive selection appears to be weak within hosts, e.g. known antigenic escape mutants do not appear to be enriched in vaccinated individuals [14, 11]. Purifying selection appears to occur to some extent. Genetic drift is thought to be strong in these populations, underscoring the role that stochasticity plays in shaping viral evolution within hosts. Furthermore, the size of the transmission bottleneck has been estimated to be on the order of one to two virions [11]. This stringent bottleneck introduces an additional source of genetic drift at the point of transmission. However, the resolution of these studies is inherently limited by the low levels of genetic diversity that exist within acute human infections of influenza [14, 11].

Analyses based solely on iSNVs do not consider the *genomic* diversity that is generated during infection in the form of defective viral genomes. These genomes feature large internal deletions in at least one segment and are therefore incapable of replicating without coinfection of a cell already harboring a wild type virus. Due to this reliance on coinfection and the spatial structure of within-host infections that may retain linkage between specific wild-type genotypes and their corresponding DVG species, we expect the ecological and evolutionary dynamics of DVGs to mirror those of the wild-type viral population.

At present, little is known about the *in vivo* dynamics of influenza DVGs. The vast majority of our understanding of influenza DVGs come from *in vitro* studies [44, 42, 17] and existing *in vivo* studies offer only a cross sectional view of DVGs within a host population [23]. How DVG populations change over time, and what those dynamics tell us about the forces shaping the entire collection of influenza viral particles within and between hosts in natural human infections therefore remained an open question. Here, we attempted to address this knowledge gap by identifying DVGs from deep sequencing data collected as part of a longitudinal influenza household cohort study [11]. We identified at least some quality-filtered DVGs in all samples in the dataset, primarily on the PB2, PB1, PA, and NS segments.

Our primary interests in this study were in assessing changes in DVG populations over the course of infection and across transmission pairs. We did not find temporal trends in the number of unique DVGs identified in an individual, nor did we find that DVG abundance (as quantified by total relative read support) changed systematically over the course of an individual’s infection. The stability in the number of unique DVG species over time implies that either DVG species persist across time or that DVGs are continually being lost and regenerated at similar rates. We found evidence for the continual loss and regeneration of DVGs, with the rate of loss of a DVG species depending on the DVG’s abundance, as measured by total relative read support.

The strength of genetic drift acting on a given population can be quantified by the effective population size *N_E_*. Drift is stronger in populations with small *N_E_* whereas the ability of stochasticity to considerably affect evolutionary dynamics will be minimal in populations with large *N_E_*. Quantifying the within-host *N_E_* of influenza A using genetic data is difficult, given the rapidly changing population sizes and noise in iSNV frequency estimates. When it has been attempted, the estimates tend to be on the order of *<* 100 [45]. While here we do not attempt to quantify *N_E_*, our observations that DVG populations are highly dynamic and change rapidly between time points is consistent with a relatively small *N_E_* as populations with a large *N_E_* would be expected to be more stable.

As discussed above, the viral diversity which is present within hosts is also shaped by the process of transmission between hosts. The size of the transmission bottleneck can be quantified to guide our understanding of how this process shapes viral diversity. To guide our knowledge of the transmission bottleneck based on the DVGs within a sample we compared DVG populations between known transmission pairs. We find that known transmission pairs, on average, share almost no DVGs and do not share more than epidemiologically unlinked pairs. Furthermore, the ones that are shared between a donor and a recipient are not present in the donor at particularly high frequencies. These findings indicate that any DVGs which are shared are likely due to *de novo* generation of the DVG in the recipient host, rather than transmission from the donor, and underscores the existence of a tight transmission bottleneck of only a small number of viral particles. While our finding that DVGs do not appear to transmit may appear unsurprising, replicating DVGs can persist in cells *in vitro* for several weeks [46], increasing the likelihood that theoretically transmitted DVGs are able to find a wild-type helper virus sometime during the course of infection. Furthermore, it is thought that viruses may transmit not independently, but in collective infectious units of multiple viral particles, which would make it more likely that a DVG and a wild-type virus from the same collective infectious unit find the same host cell [25]. Our finding that DVGs do not appear to be transmitted from donors to recipients is consistent with the existence of a very stringent transmission bottleneck, and indicates that small transmission bottlenecks may benefit a viral population by purging viral “cheaters’’ in the form of DVGs [47].

There are some important limitations to our analyses. First, as with any *in vivo* deep sequencing based analysis, the samples taken from individuals using throat or nasal swabs may not adequately reflect the within-host viral population, particularly when sub-compartmentalization within a host is substantial as has been shown in experimentally infected animal hosts [48, 49]. Second, it is difficult to reliably quantify the amount of DVGs within a sample using sequencing data. Here we have relied on a relative measure of the number of reads spanning a given junction site to the total number of reads mapped to a given segment. This method is slightly biased, however, because we cannot assign reads from the 5’ and 3’ termini as belonging to either a wild type or DVG segment. Furthermore, this metric will be biased downwards in longer segments as there are more coding nucleotides which can be deleted in a DVG without affecting the packaging signals. However, without additional laboratory measurements, we feel this metric represents a suitable attempt to account for varying sequencing depth between samples. Furthermore, our set of identified DVGs will be affected by the amplification and sequencing protocol used to generate these data. Genetic material was amplified based on primers which bind to the terminal regions of the wild-type segments and there are several size-filtering steps which eliminate smaller DVG segments. However, there is reason to expect that DVGs without the terminal packaging signals would be unlikely to be efficiently packaged into virons, even if they were generated by the polymerase [43]. Despite these limitations, the fact that the plasmid controls harbor very low relative amounts of DVGs provides confidence that the DVGs reported here represent a subset of true biological DVGs. Alternative viral sequencing approaches, particularly long read sequencing, have the potential to overcome some of these shortcomings and in future studies may provide greater resolution into the evolutionary processes shaping within-and between-host viral populations.

## Supporting information

**S1 Fig.**
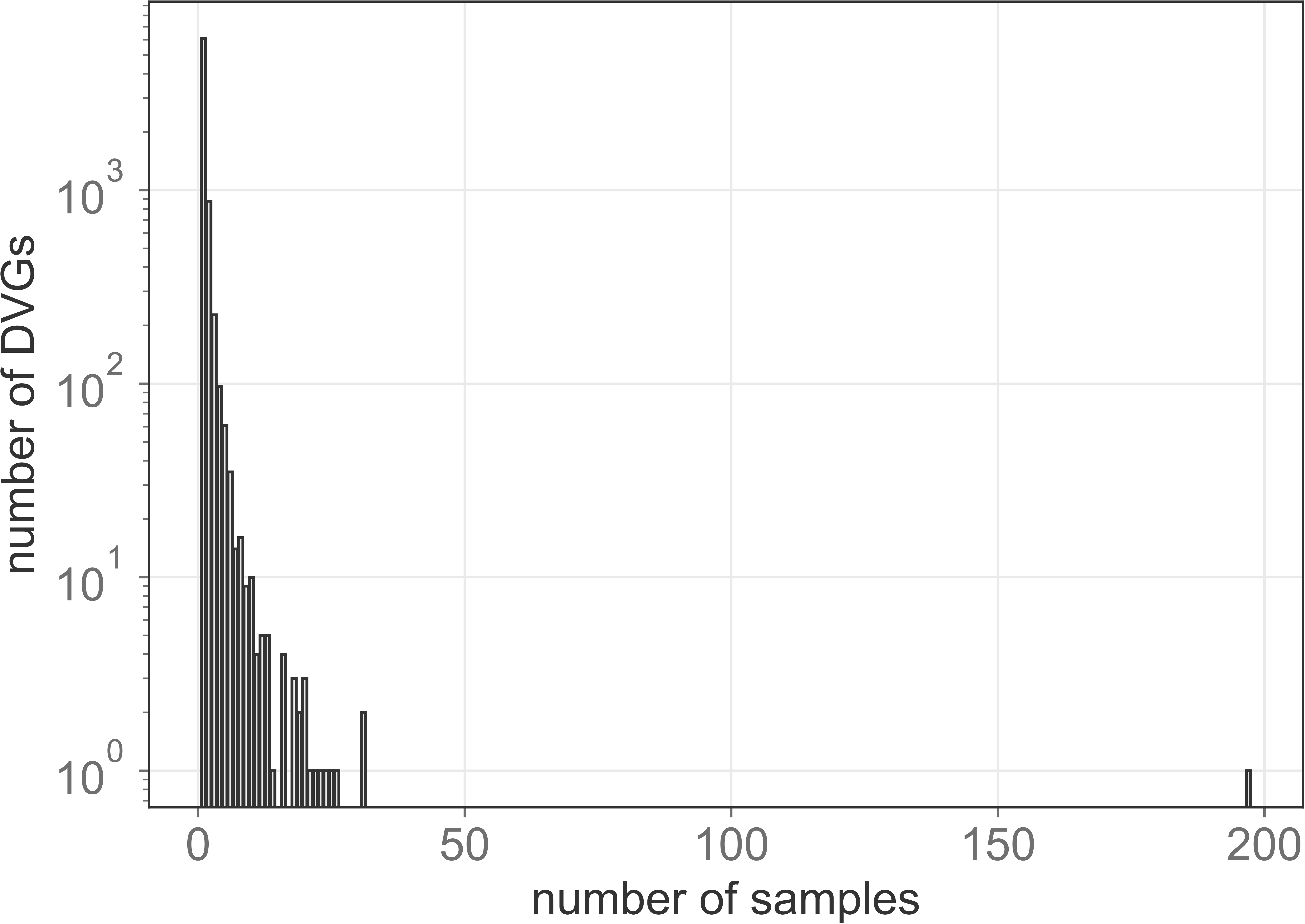
Prevalence of unique DVG species. Number of clinical samples in which each unique DVG species (as identified by the segment and breakpoints) is identified in.

**S2 Fig.**
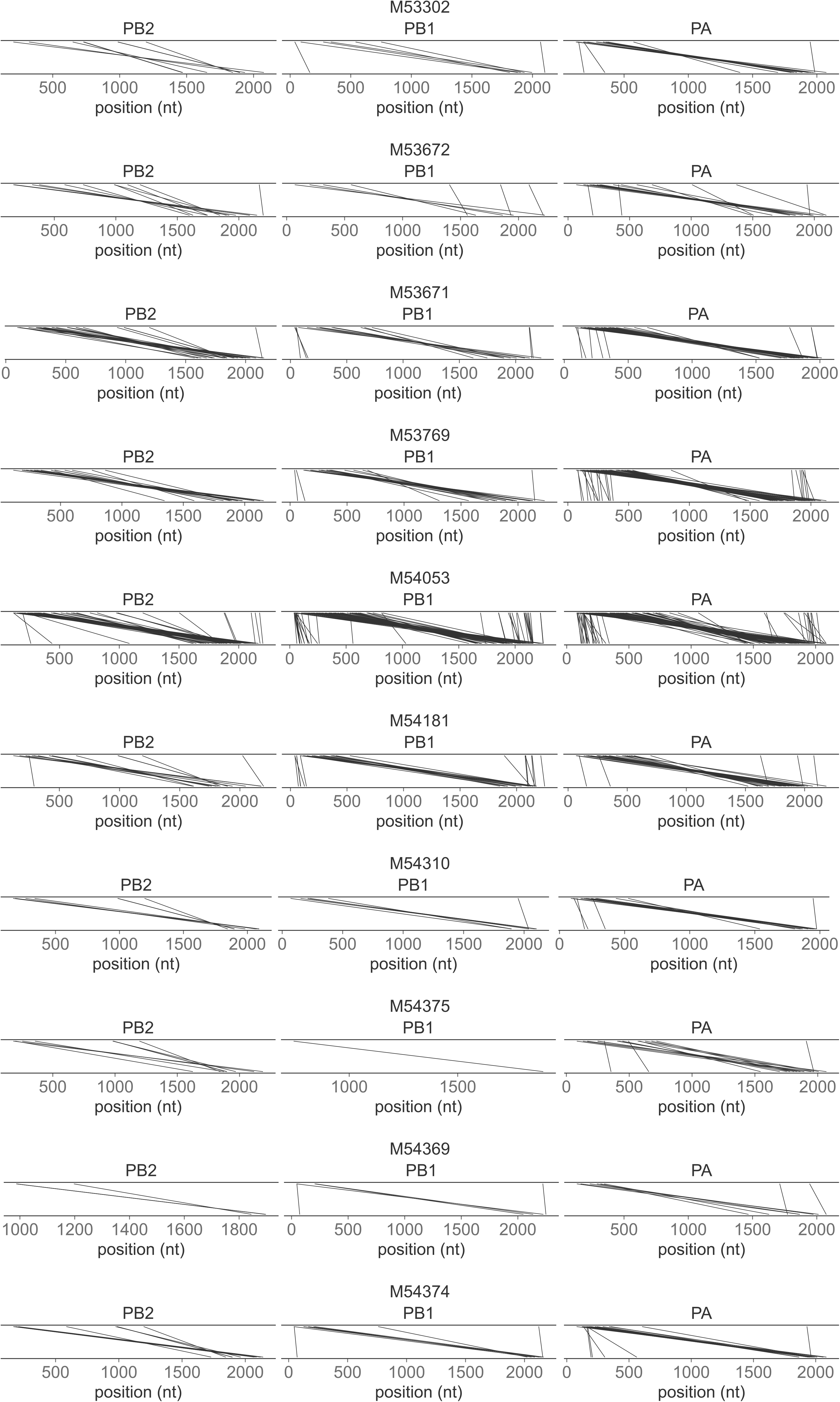
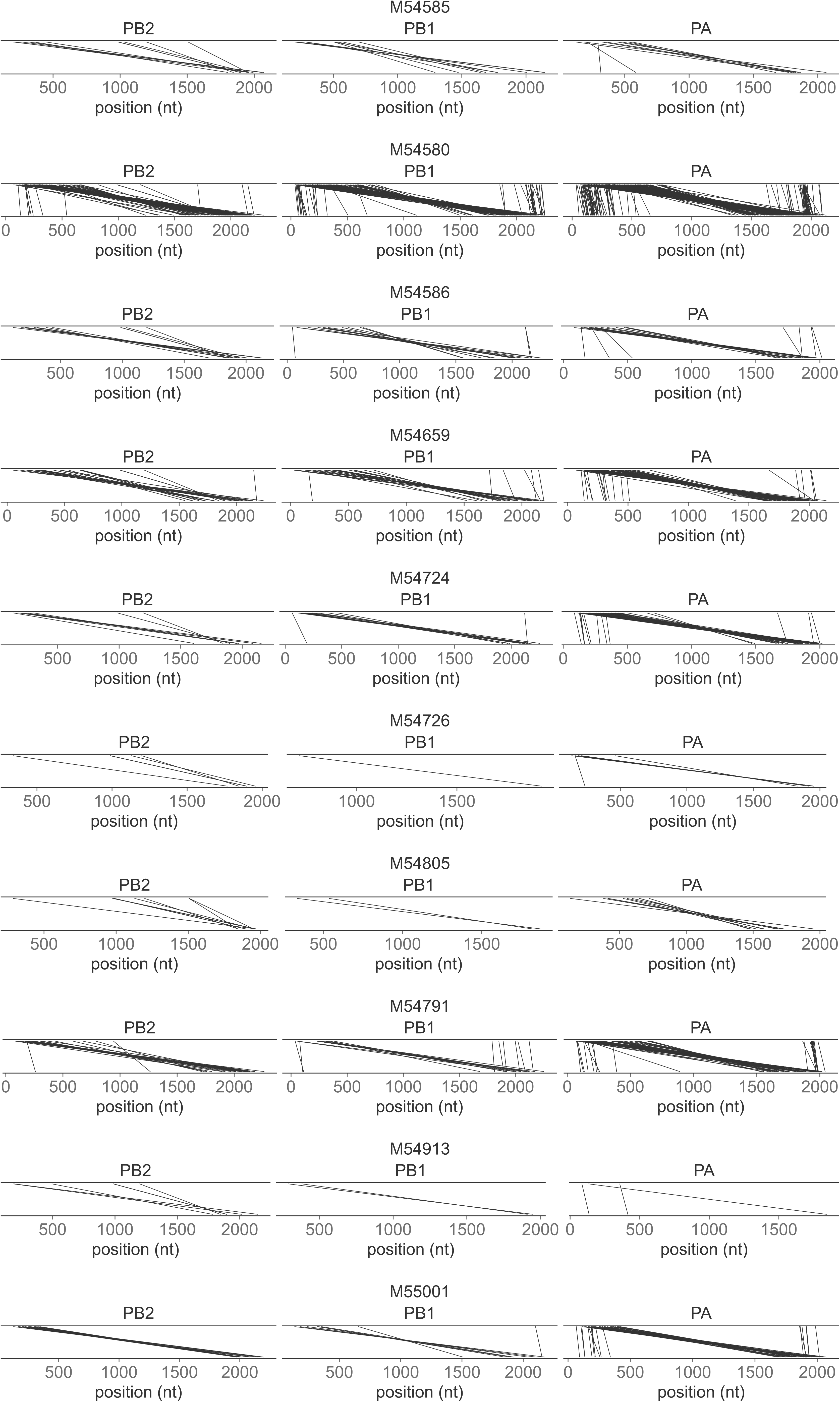
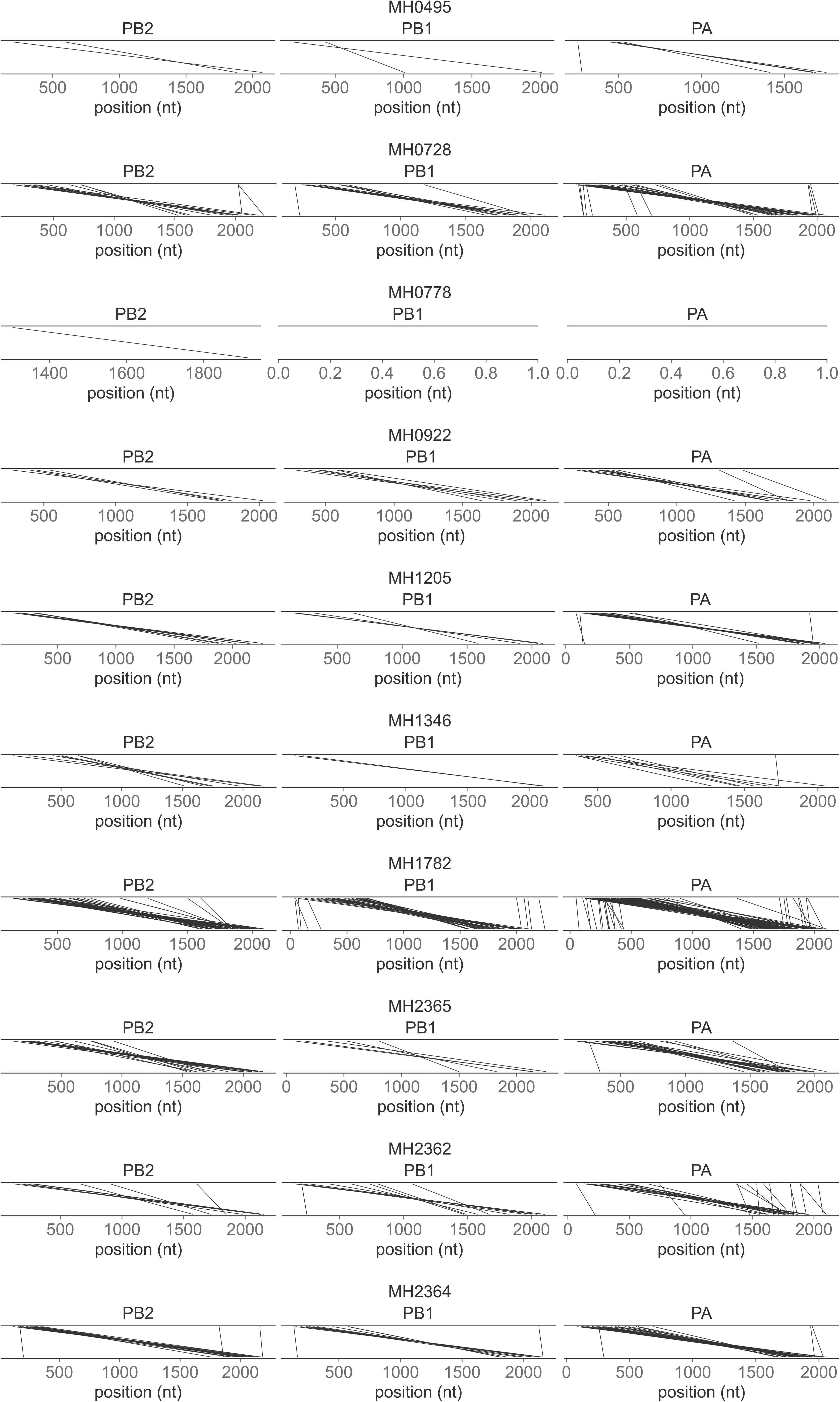
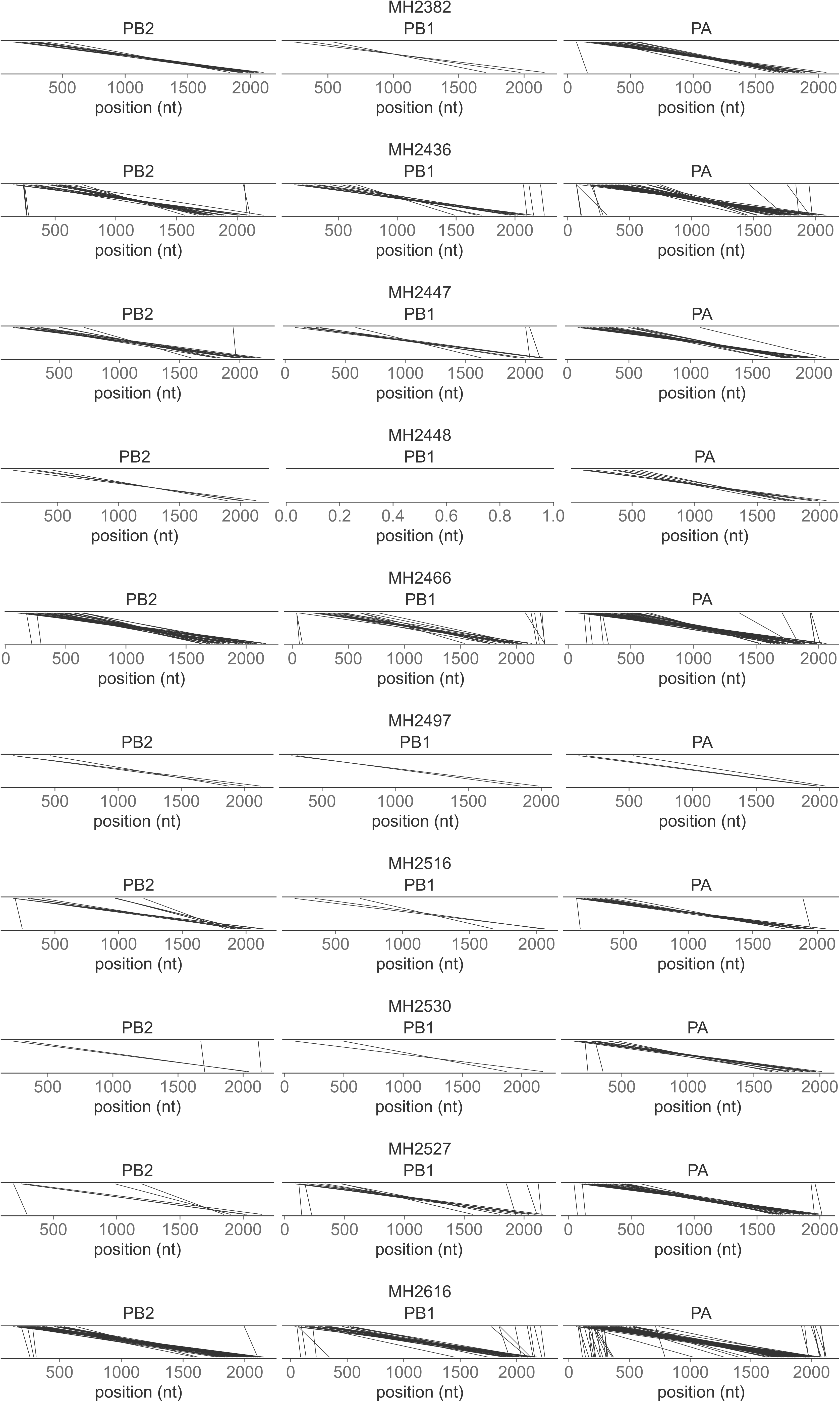
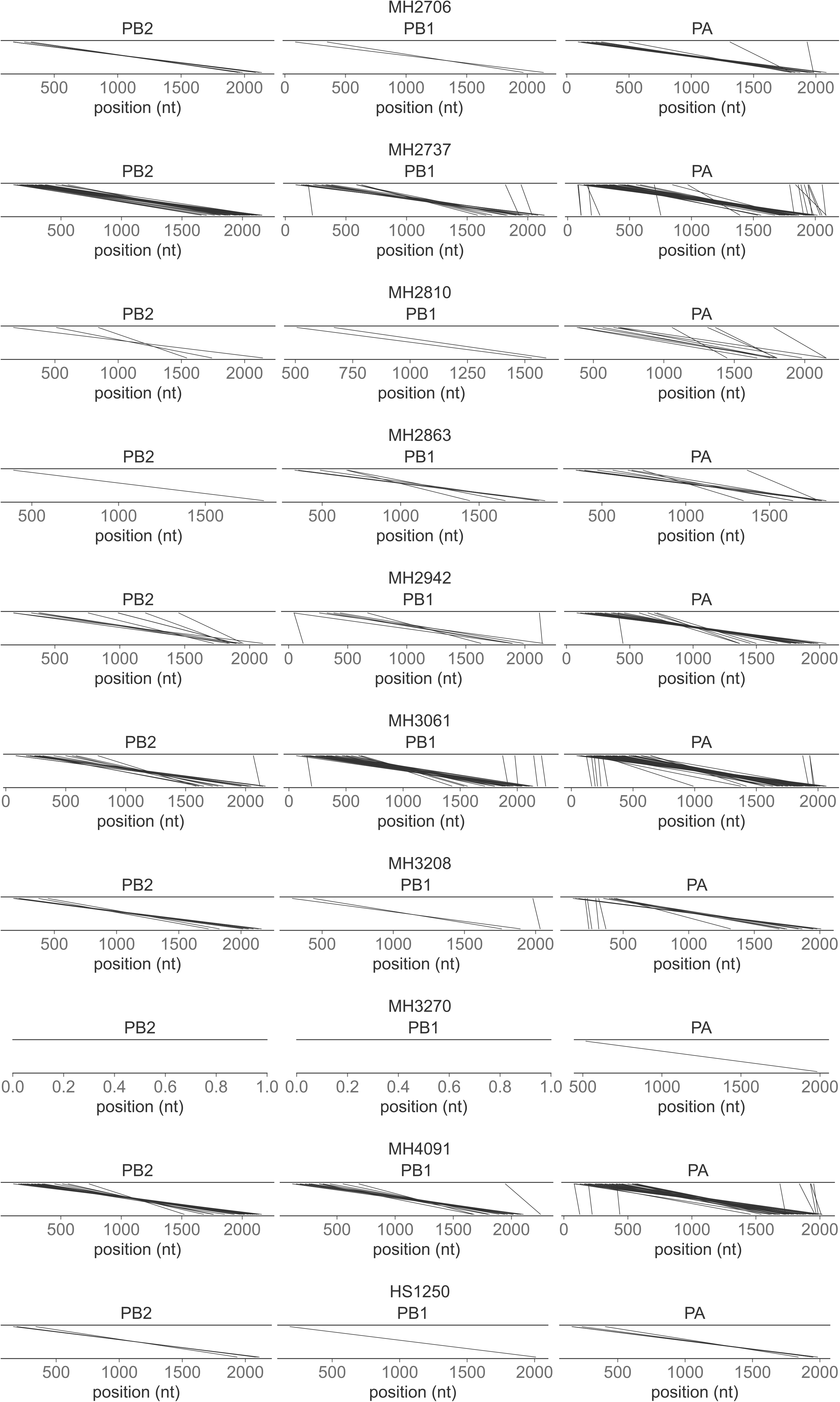
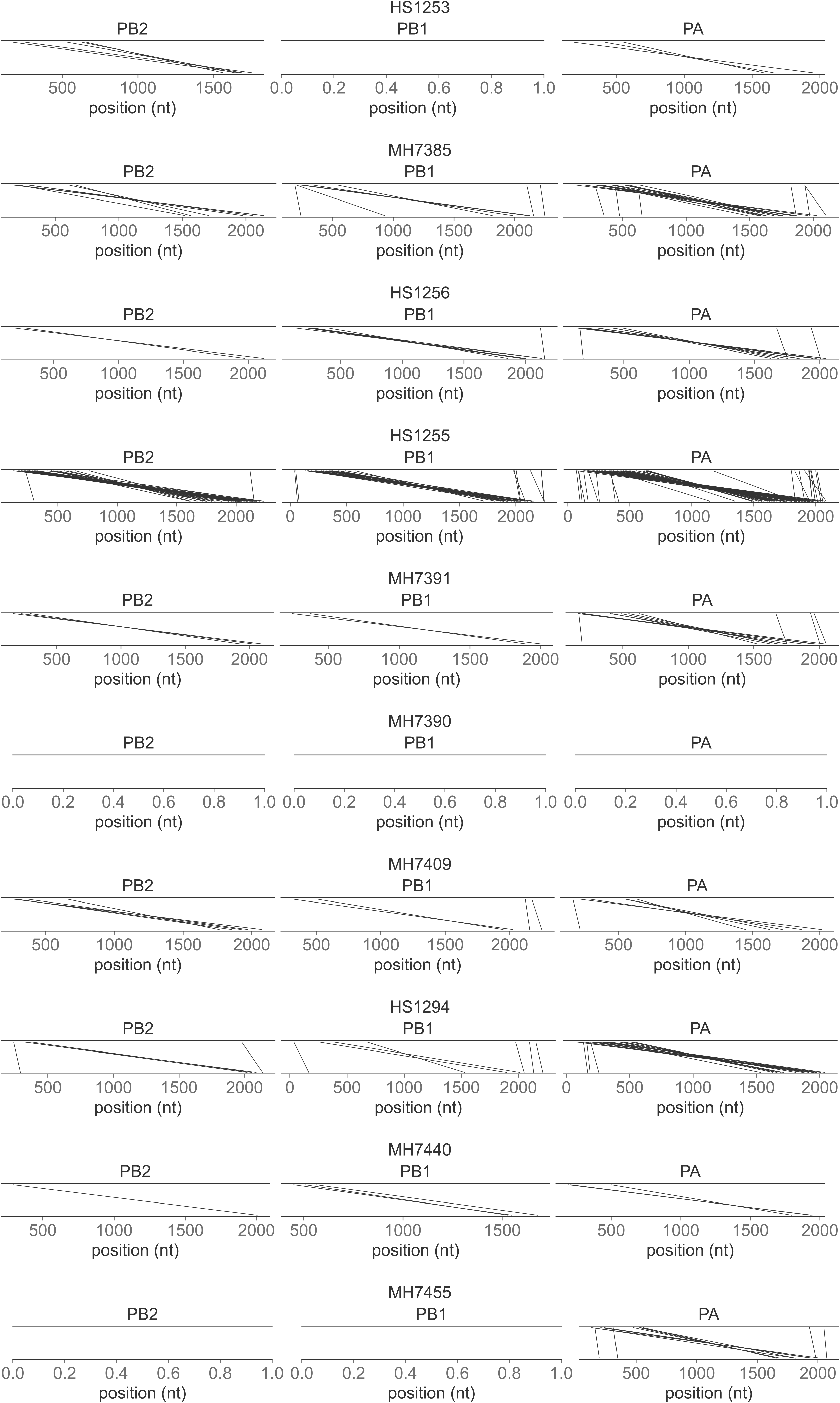
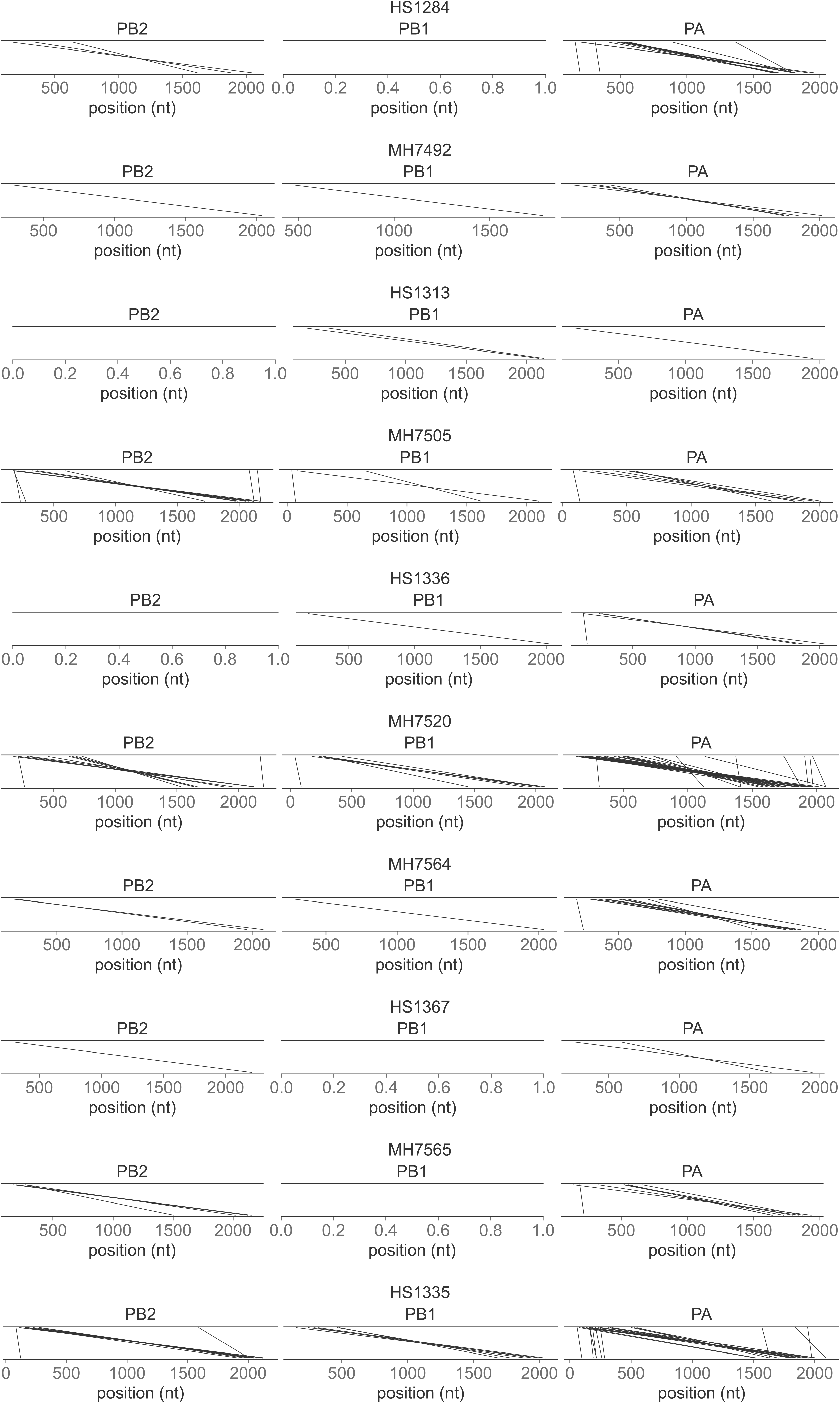
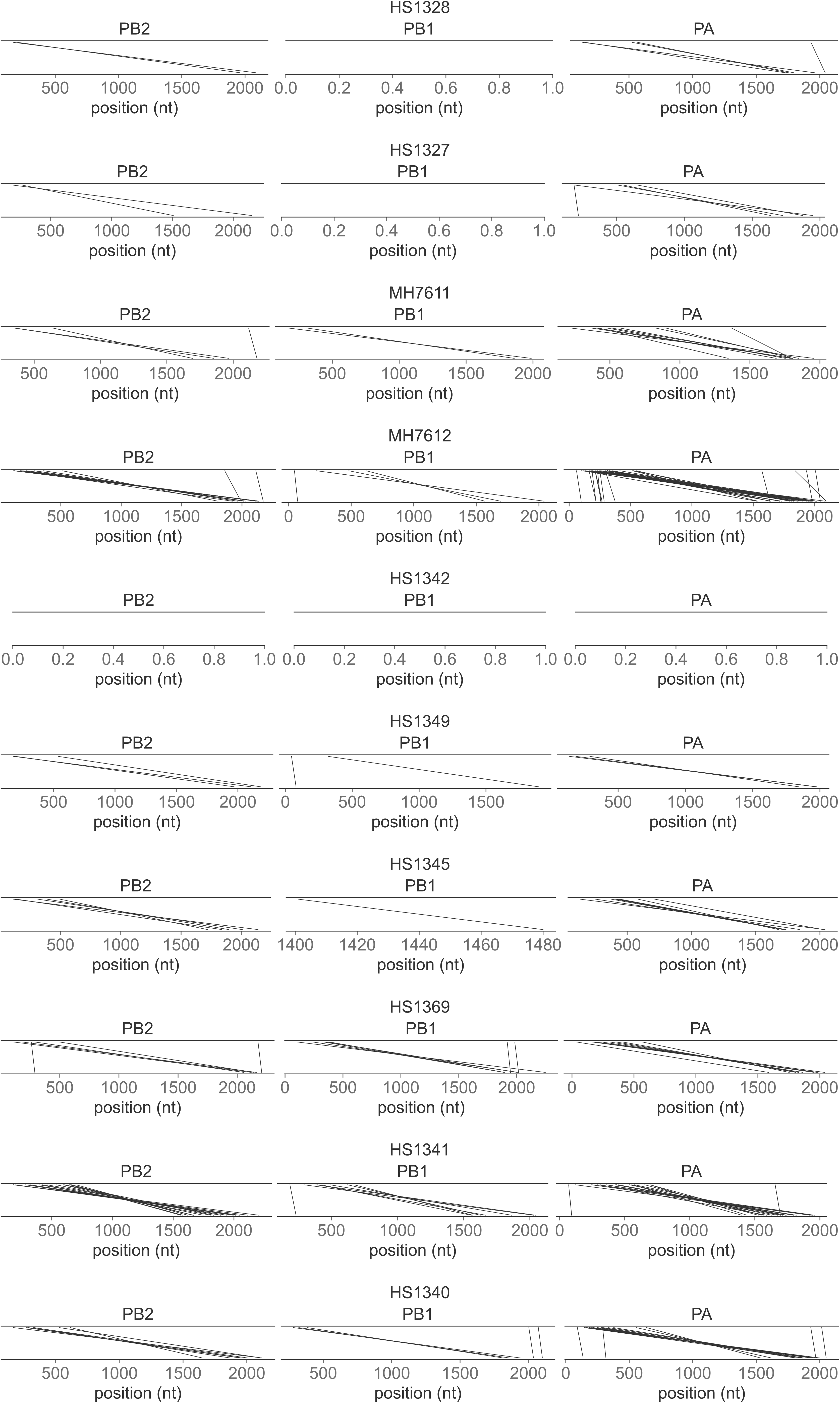
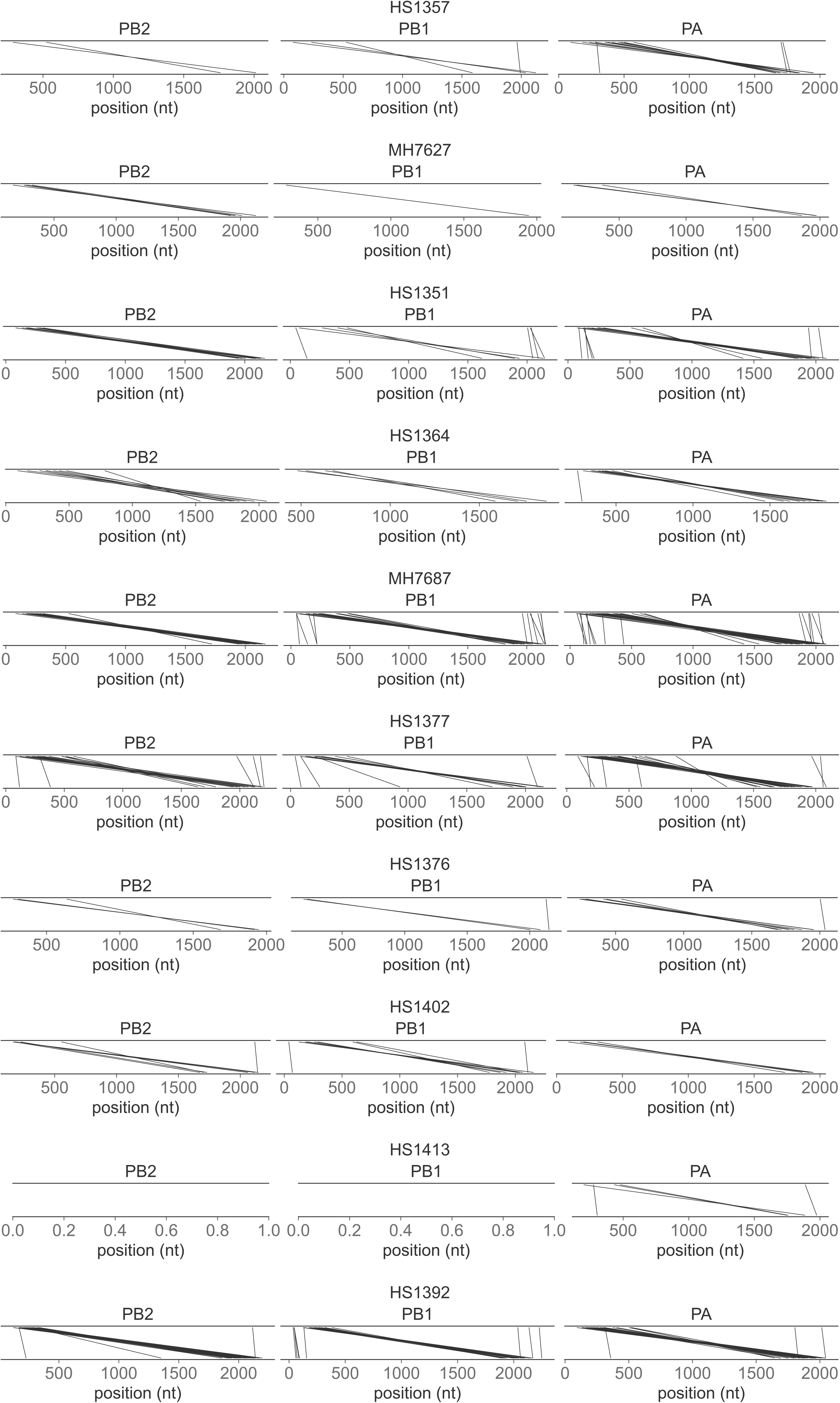
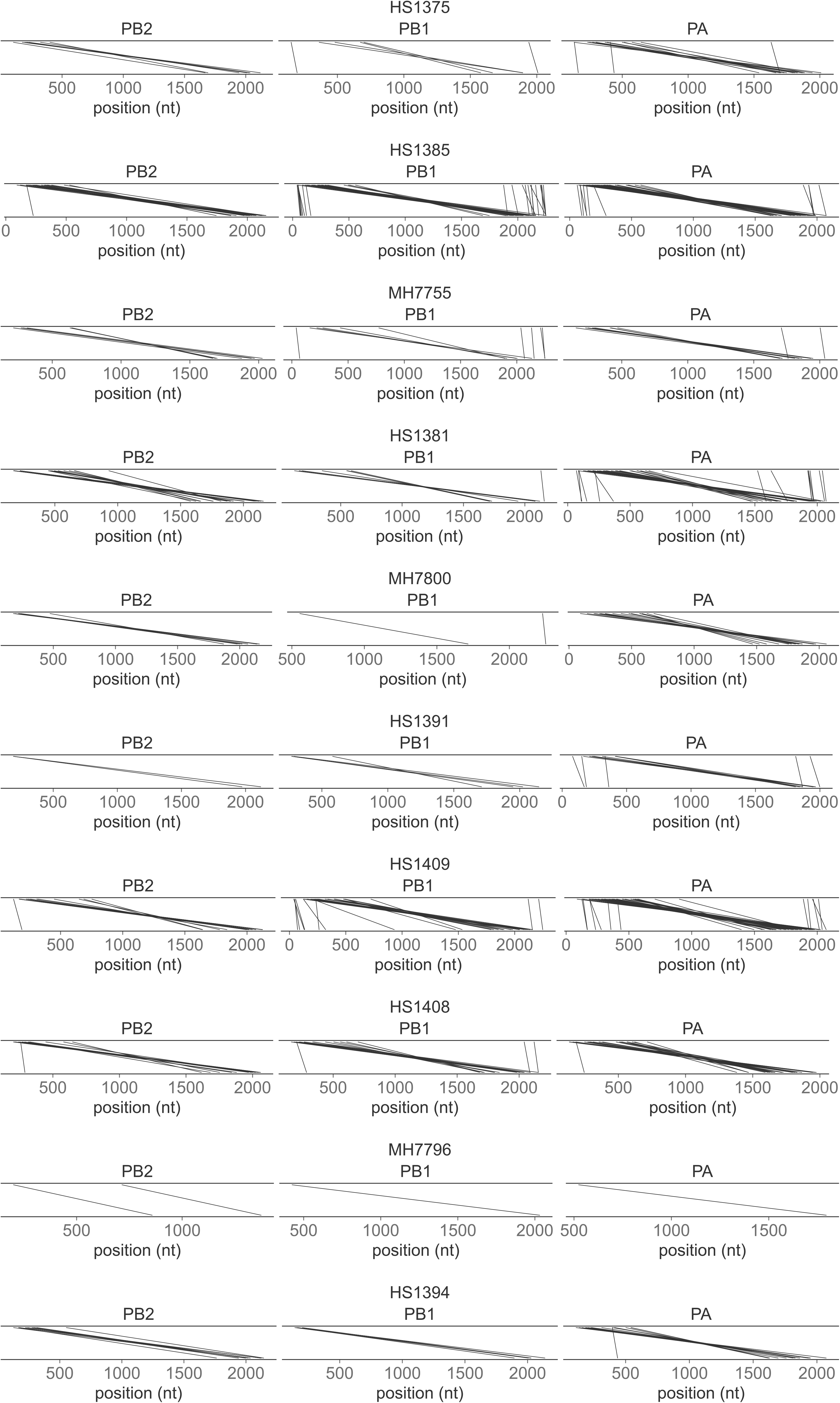
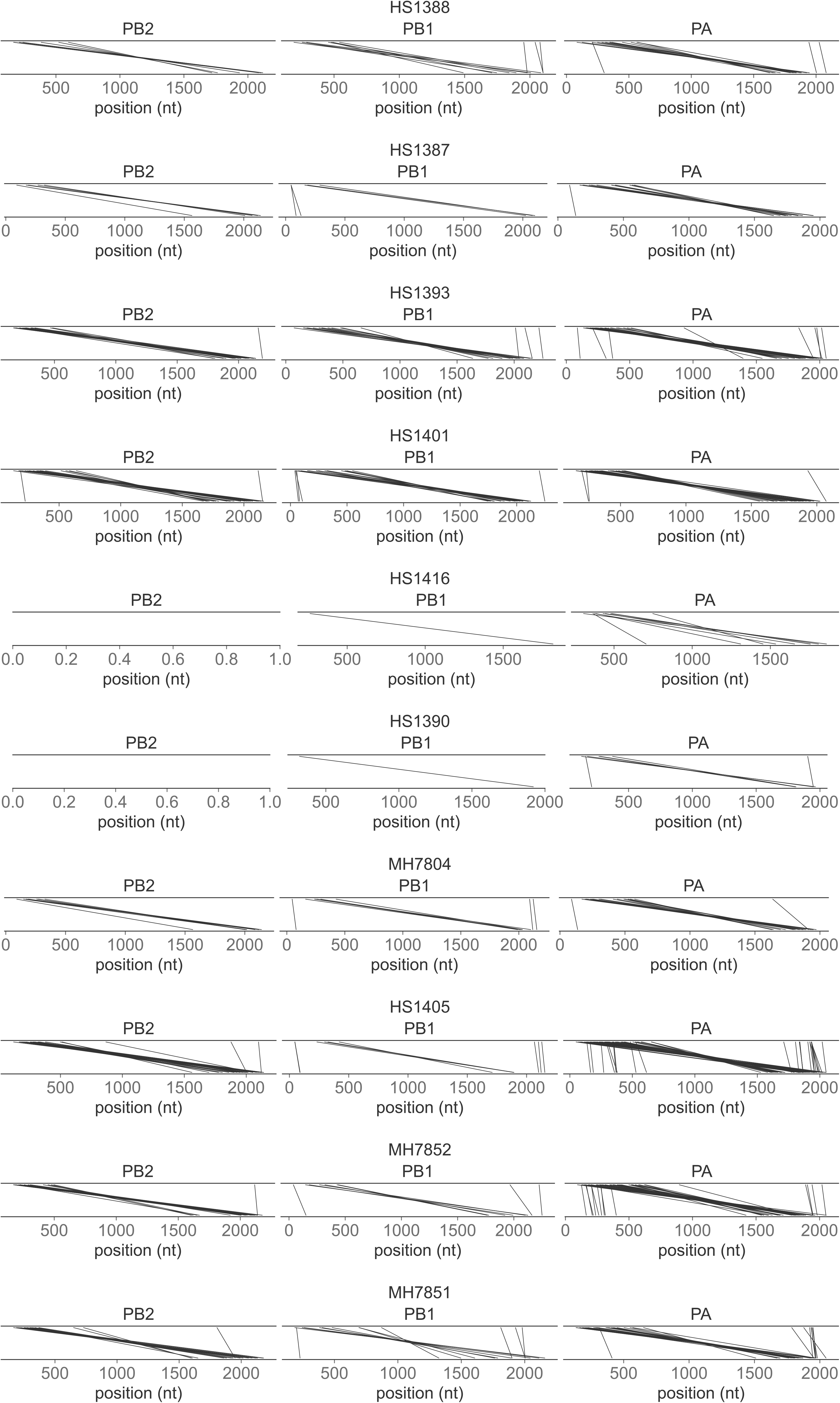
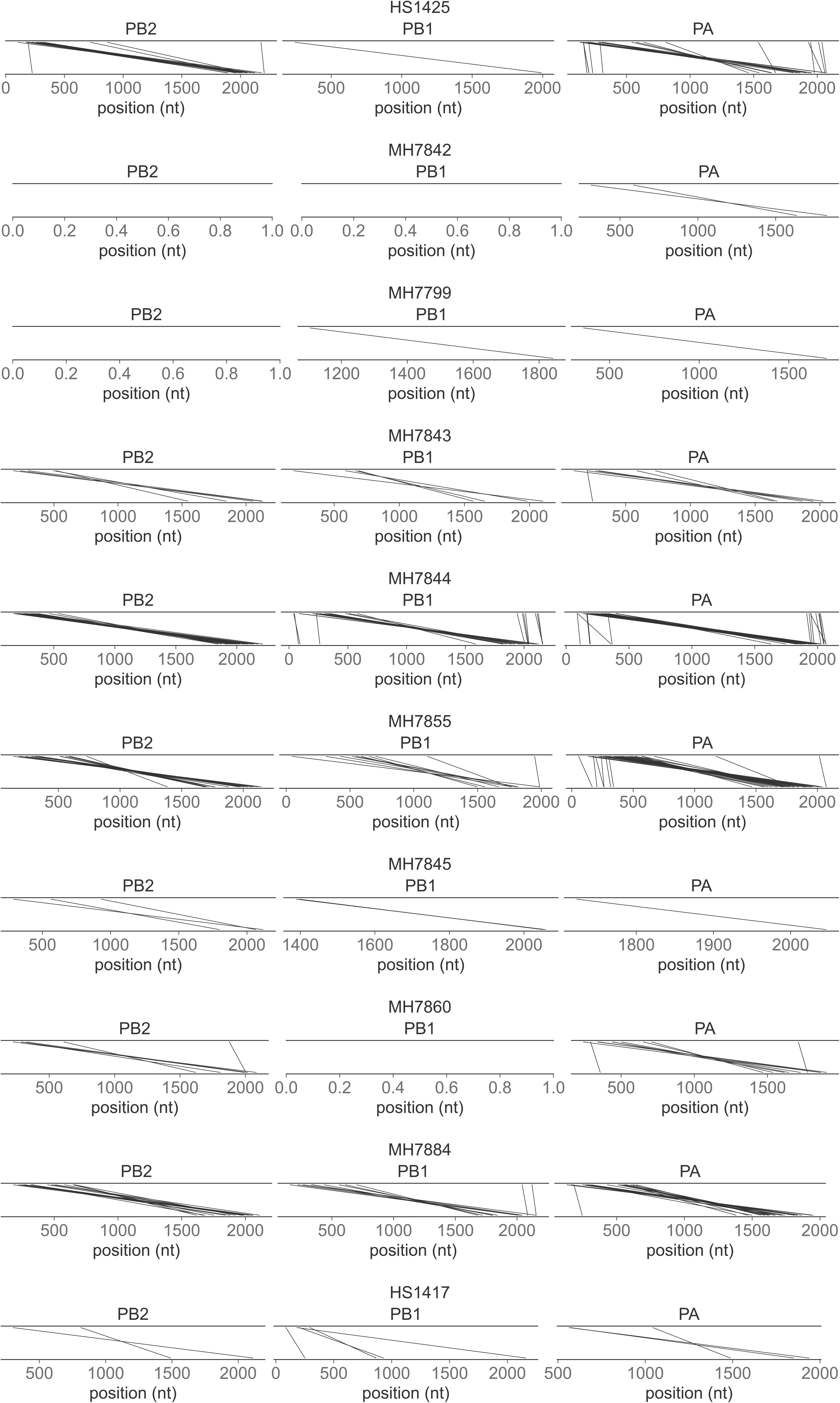
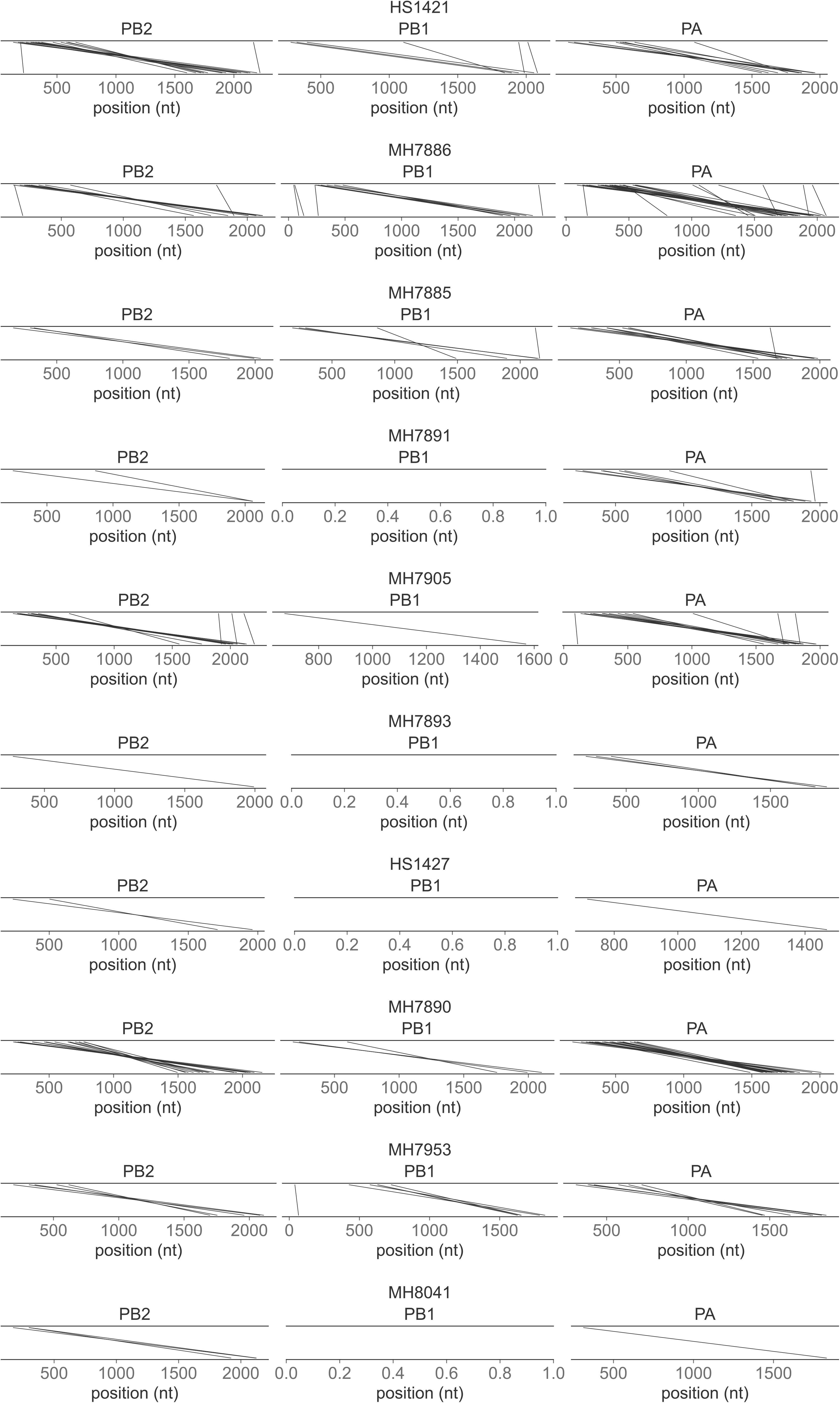
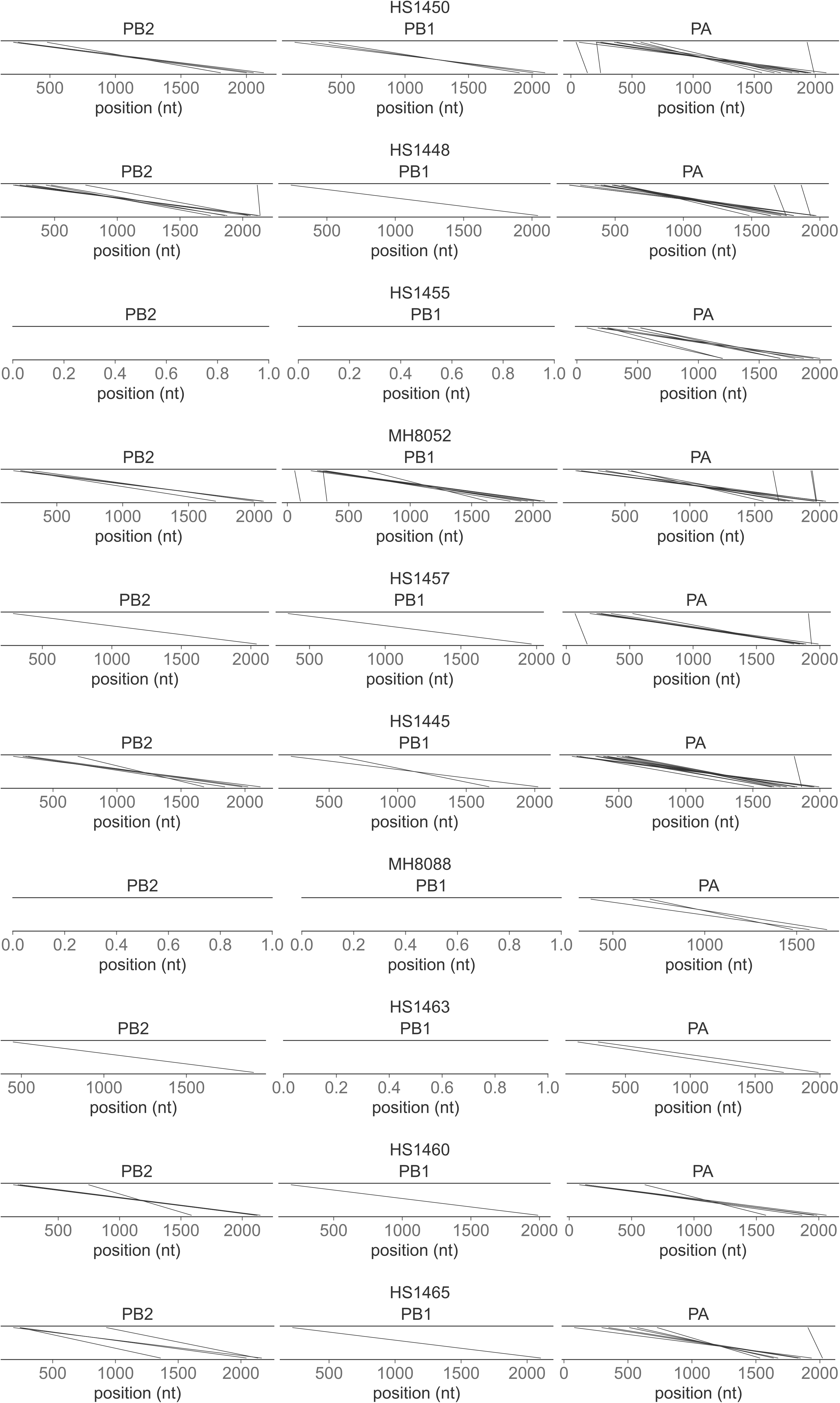
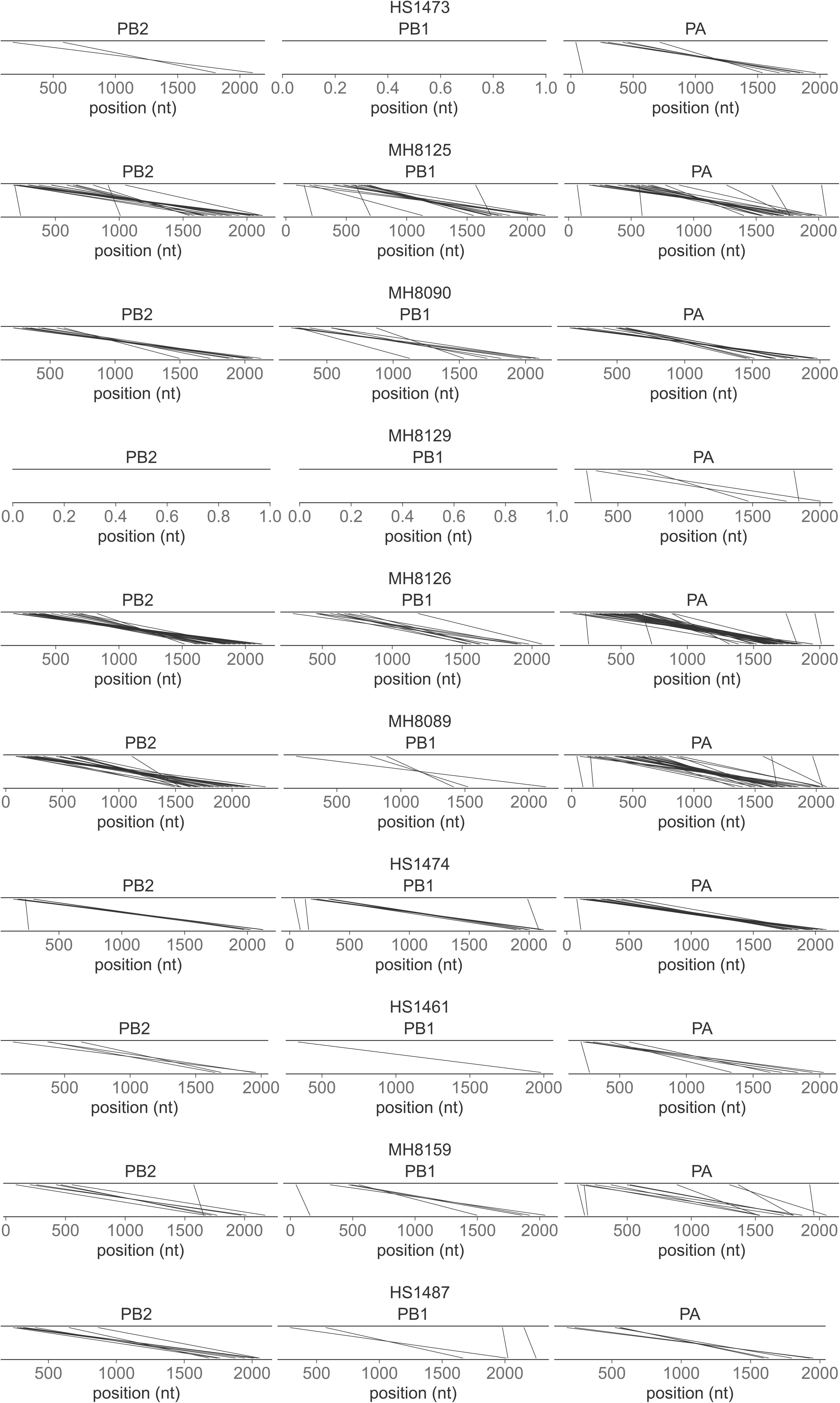
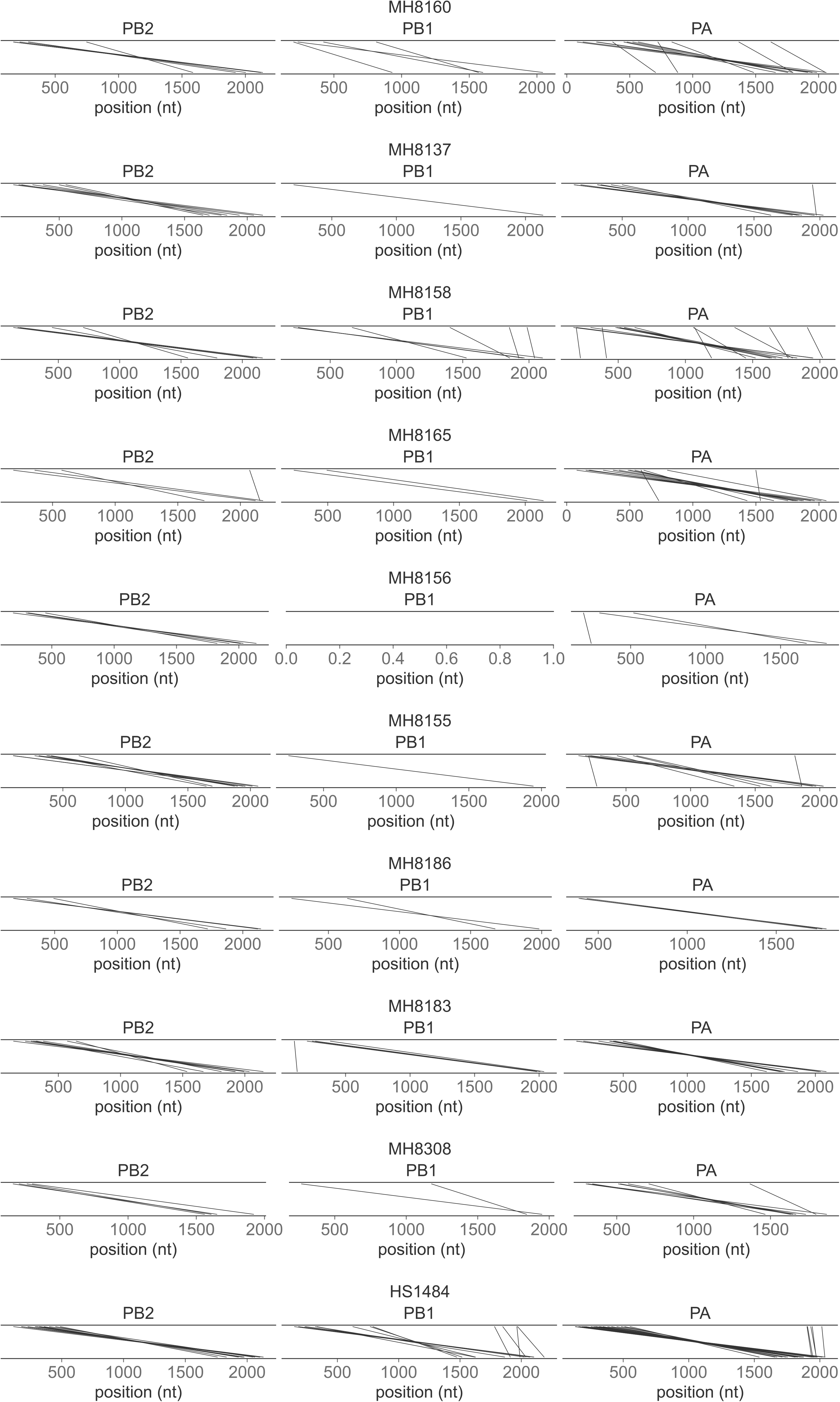
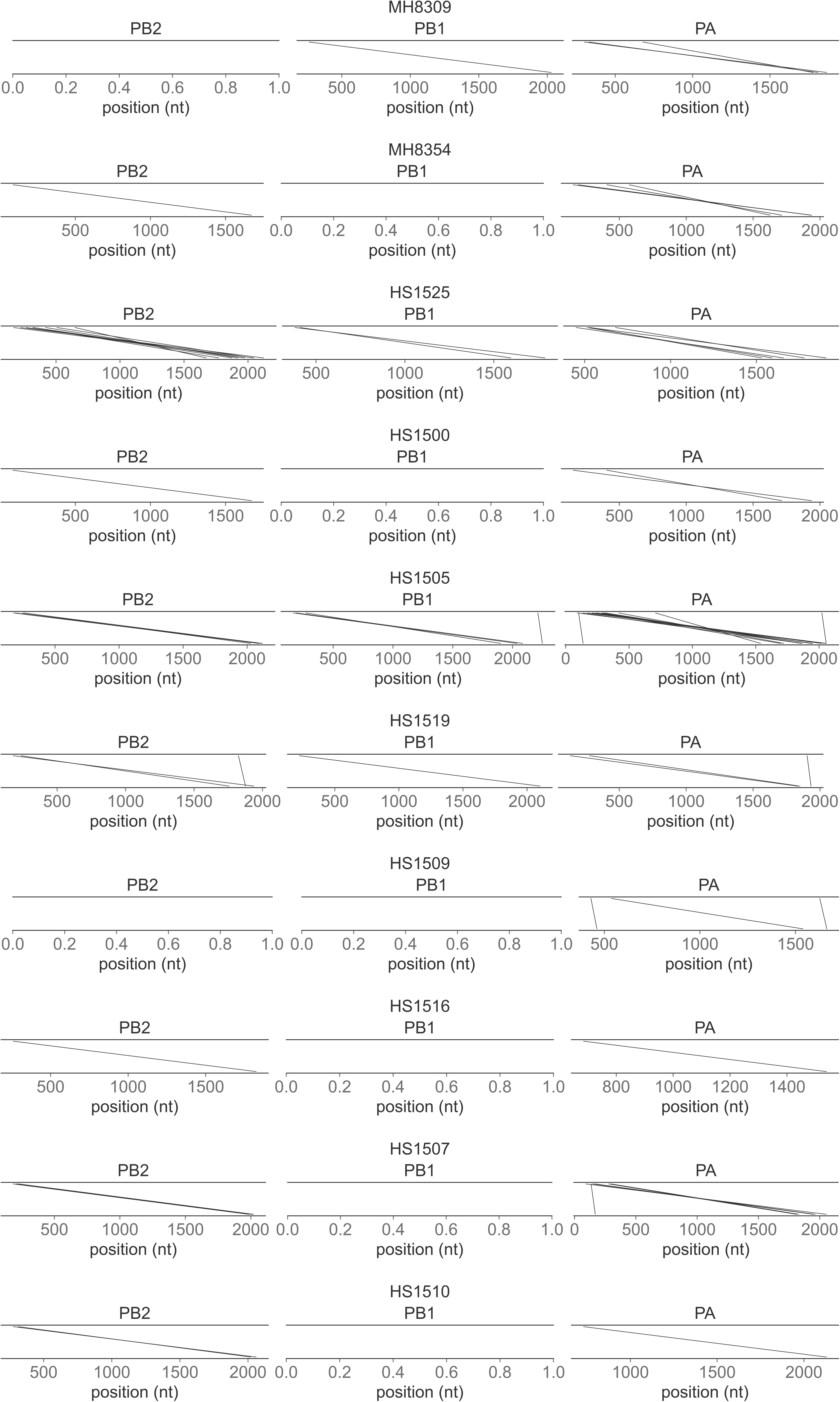
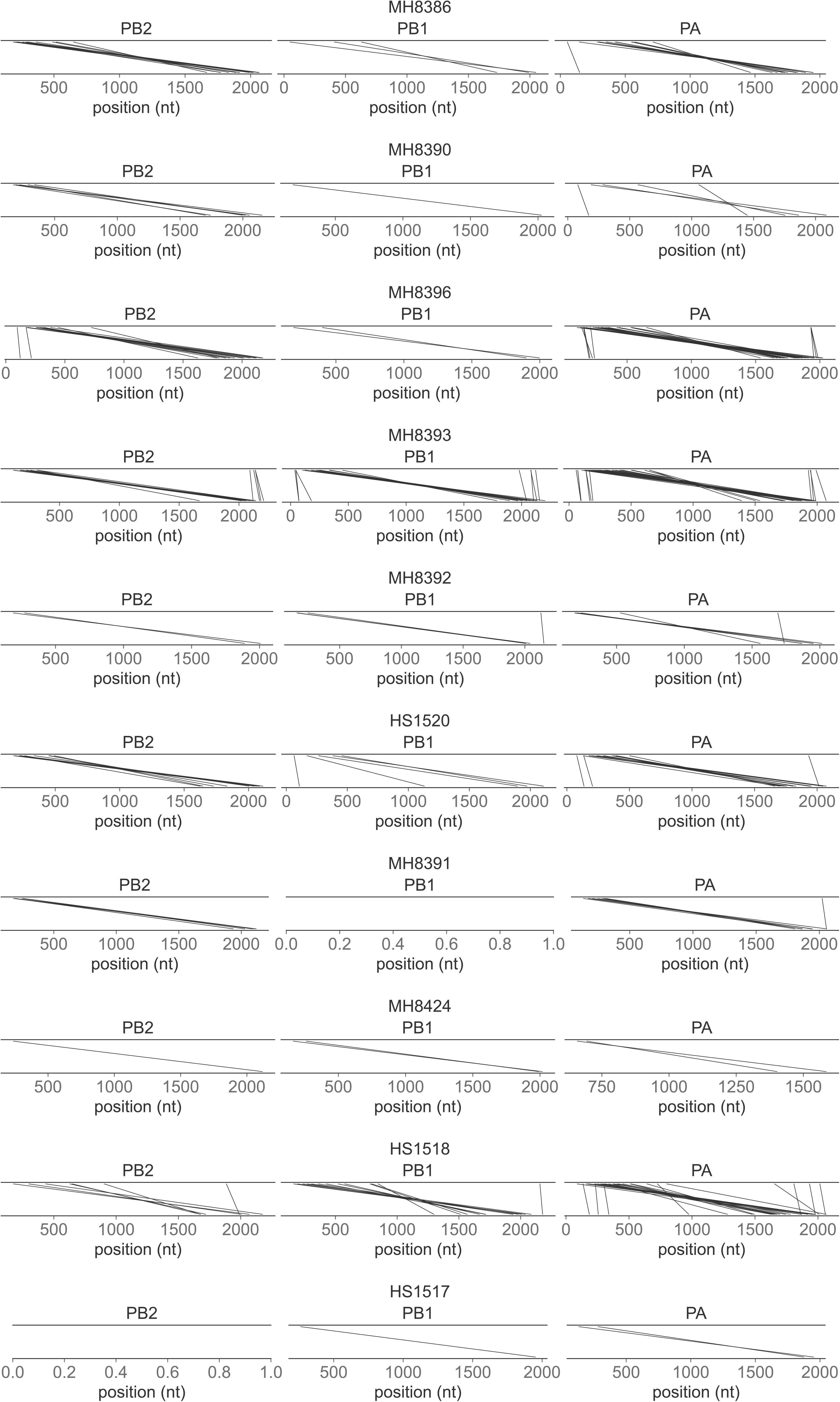
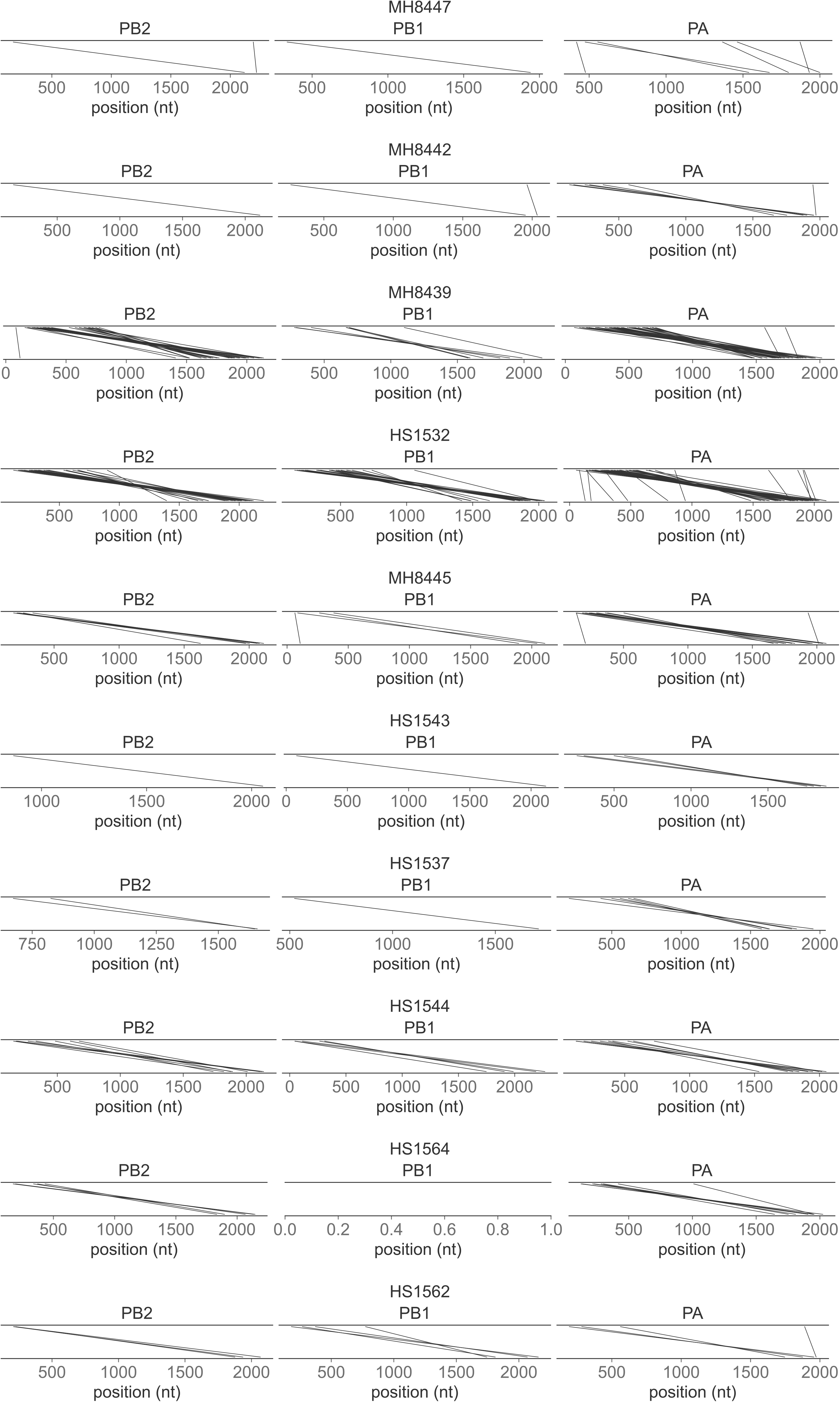
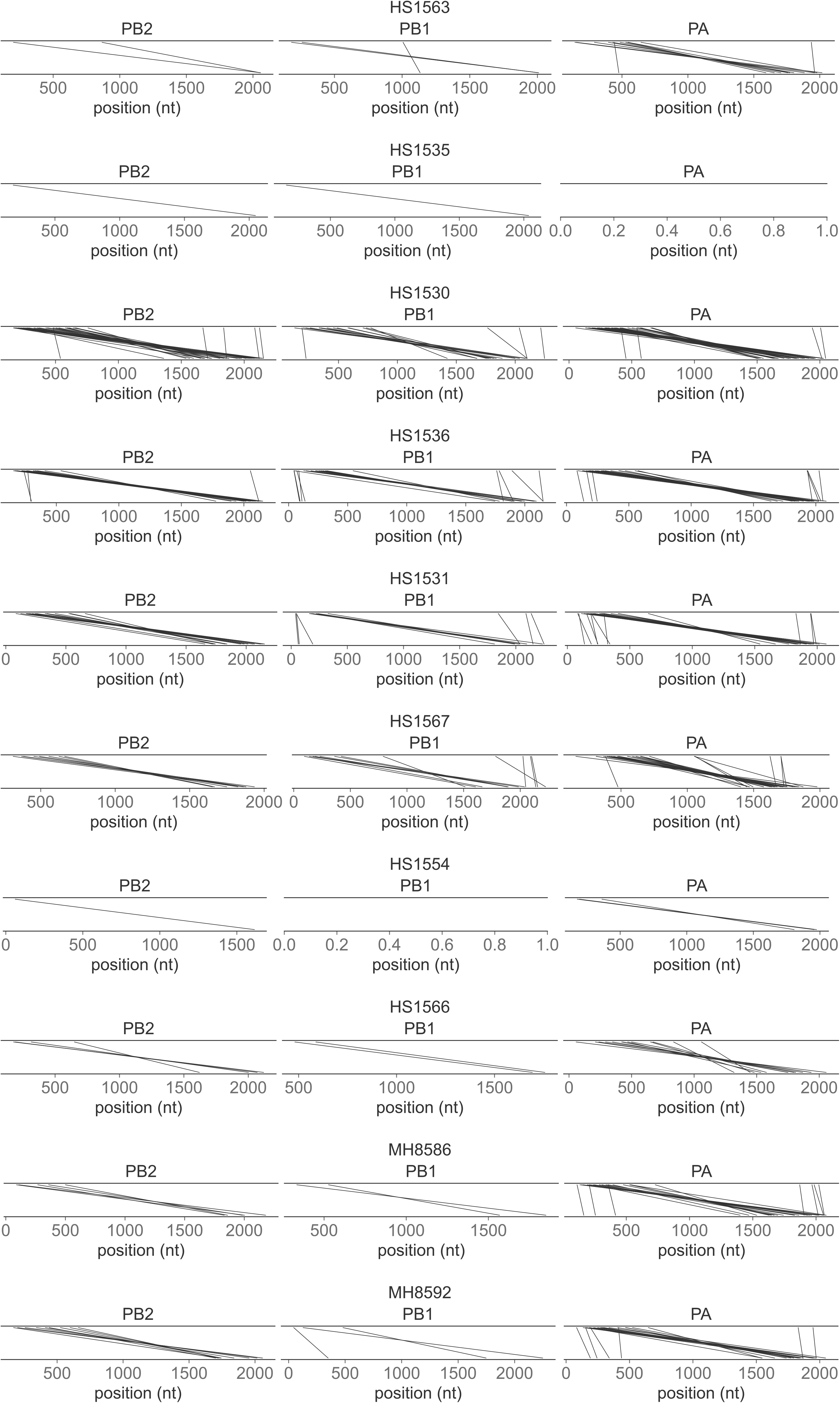
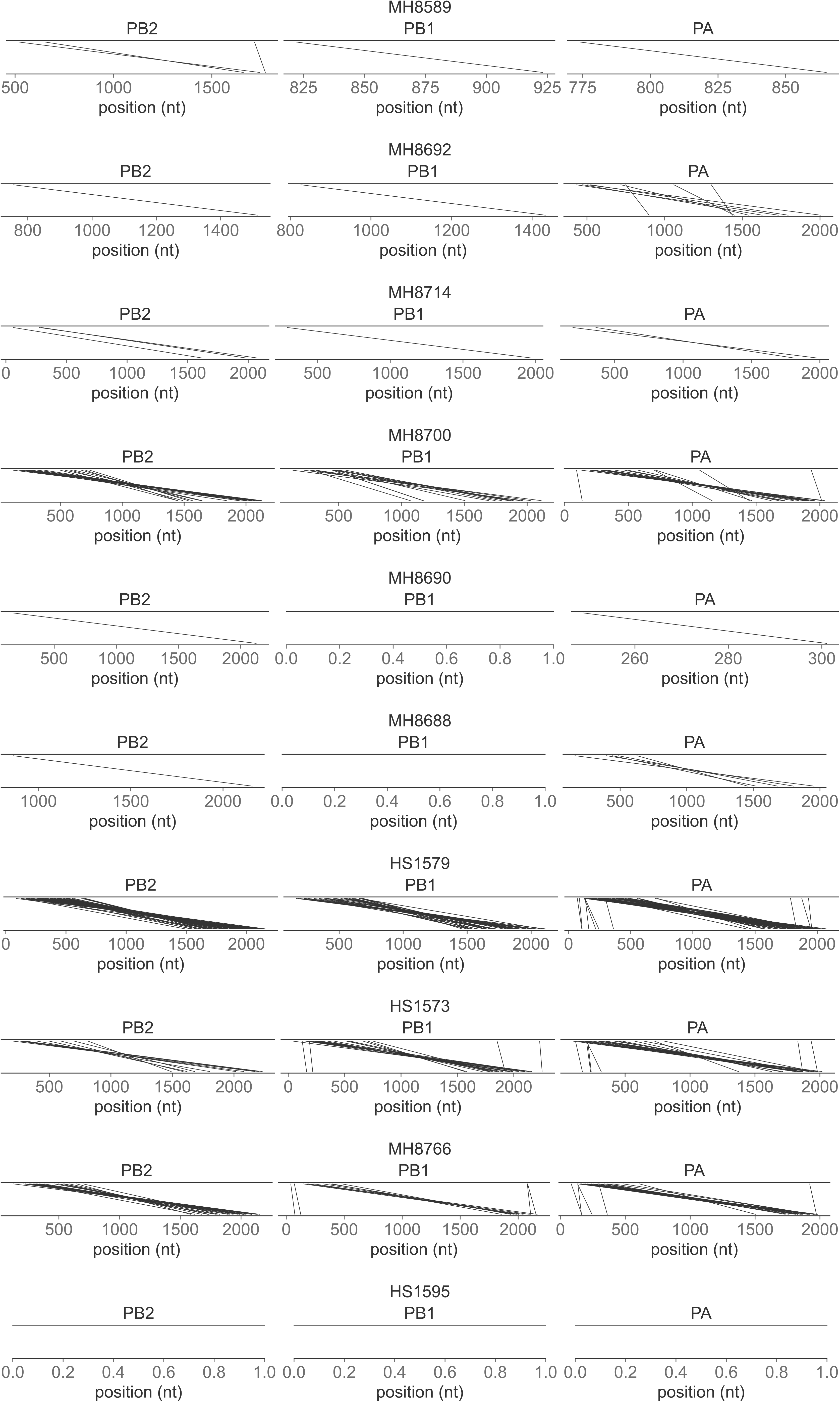
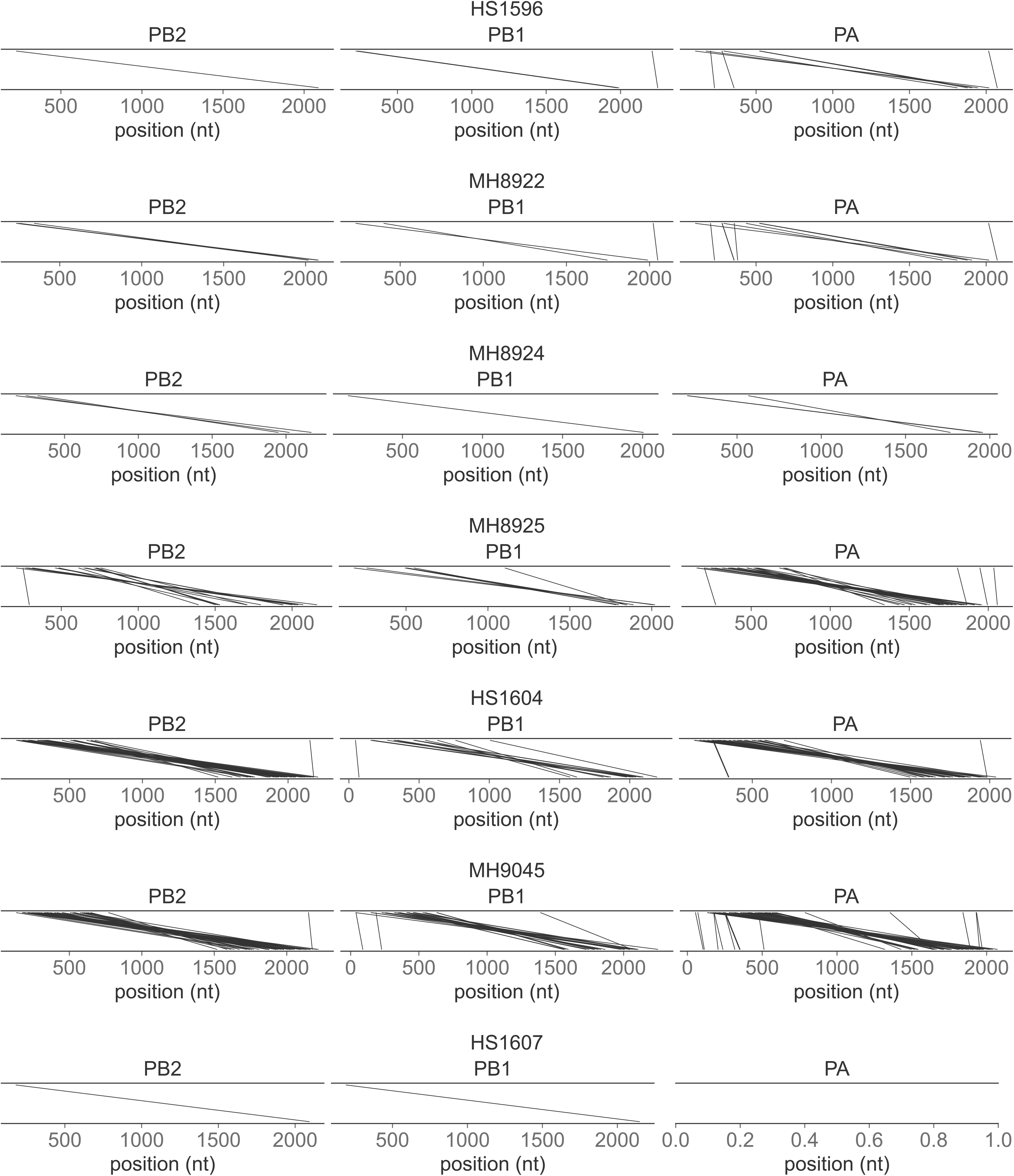
DVG junction locations. Observed junction locations for all PB2, PB1, and PA DVGs observed in all clinical samples. Lines connect the nucleotides flanking the deleted nucleotides for each identified DVG.

**S3 Fig.**
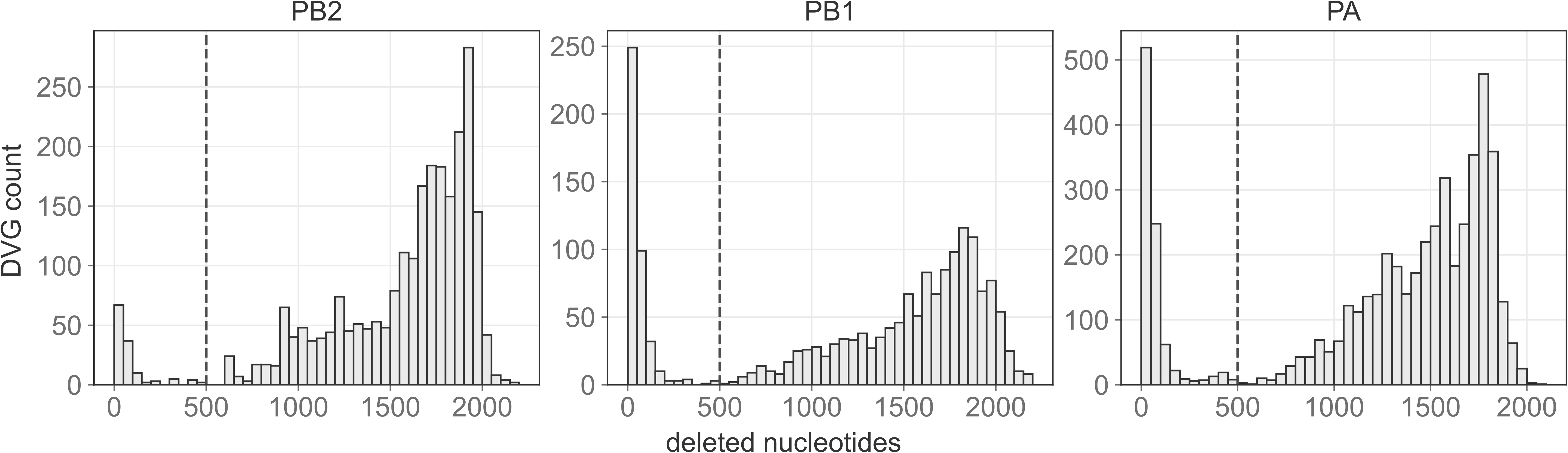
Deleted nucleotides. Number of deleted nucleotides for each DVG identified in the PB2 (A), PB1 (B), and PA (C) segments. Dashed line at 500 nucleotides represents our empirical filtering threshold.

**S4 Fig.**
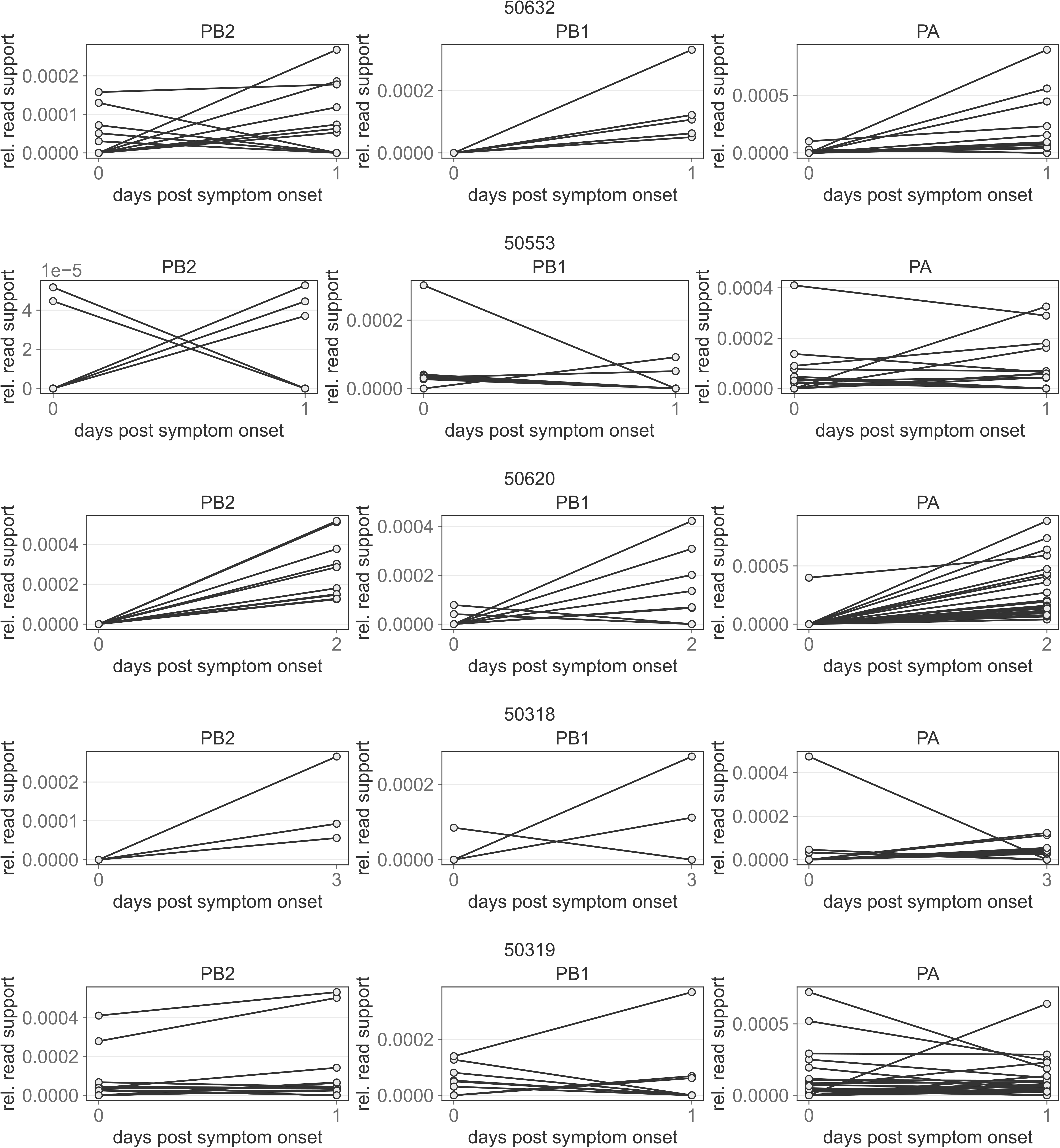
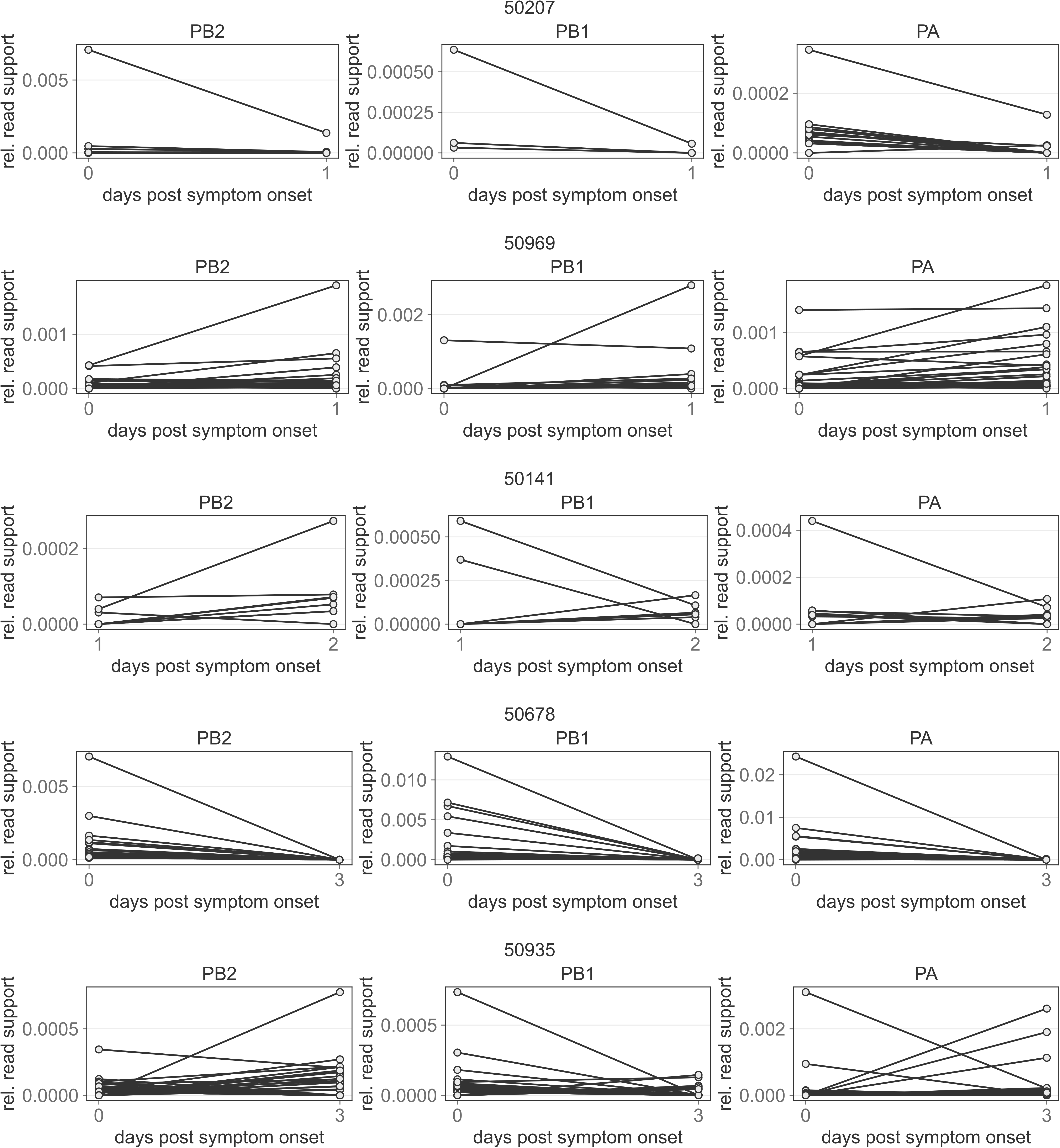
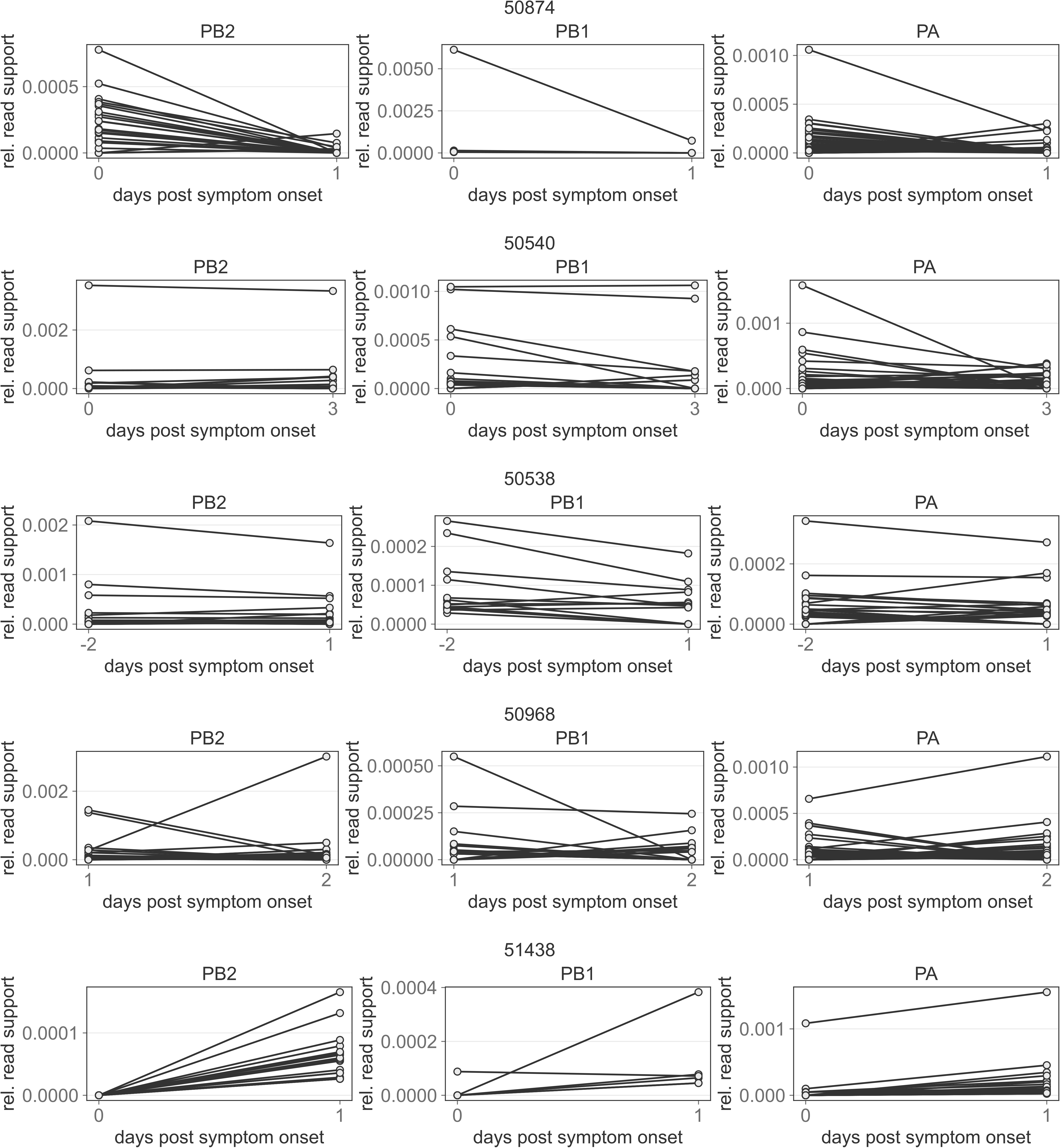
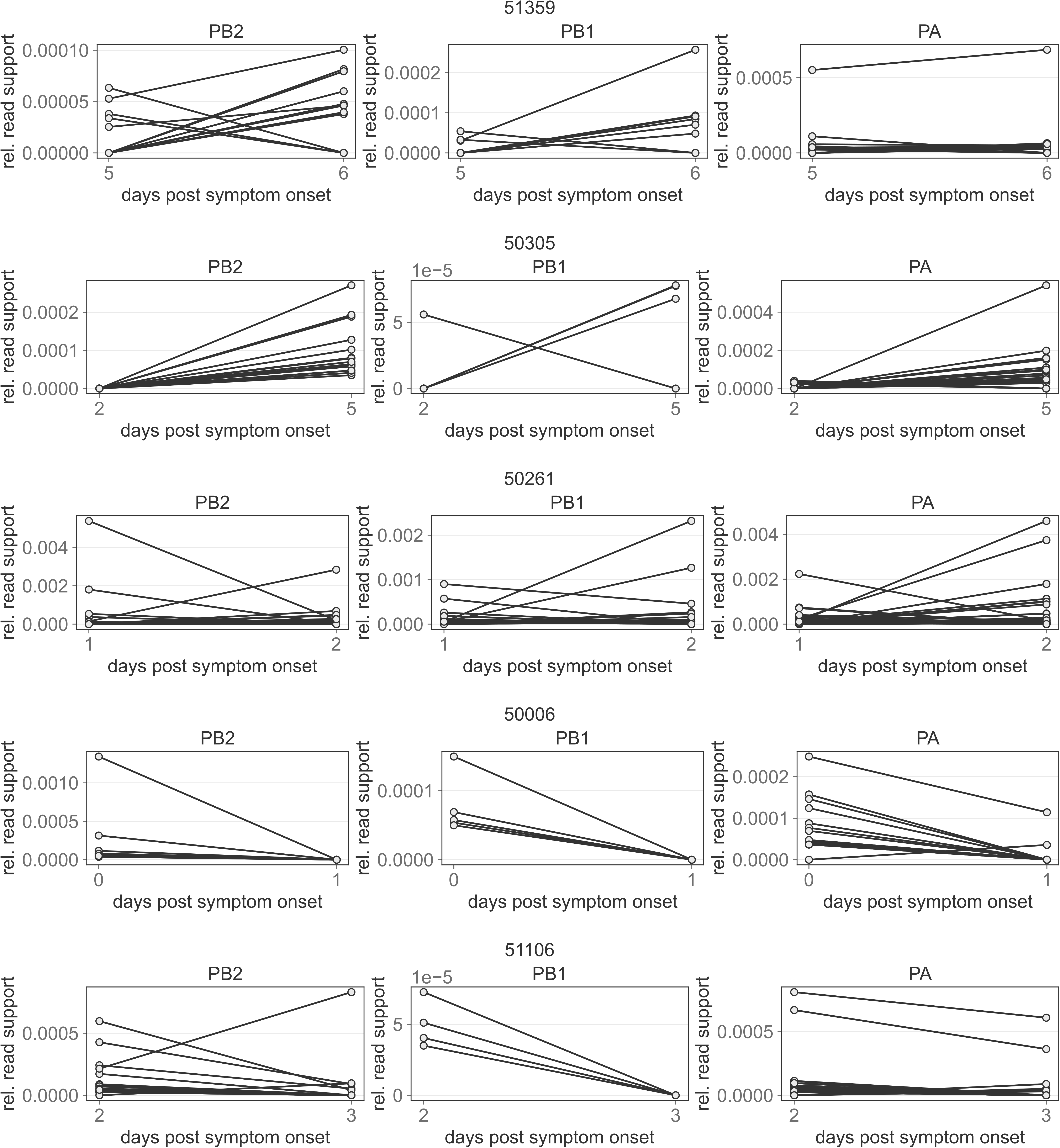
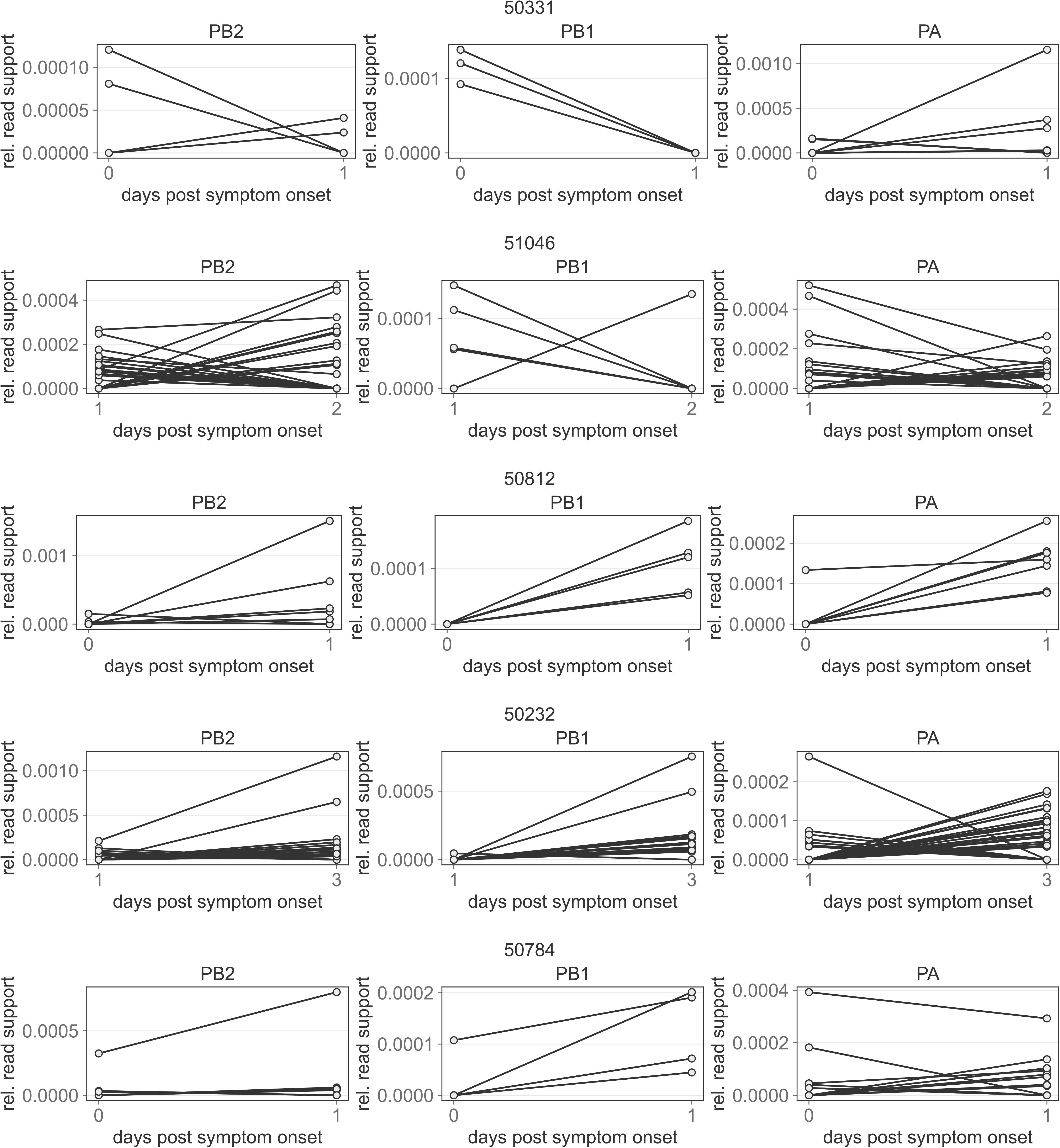
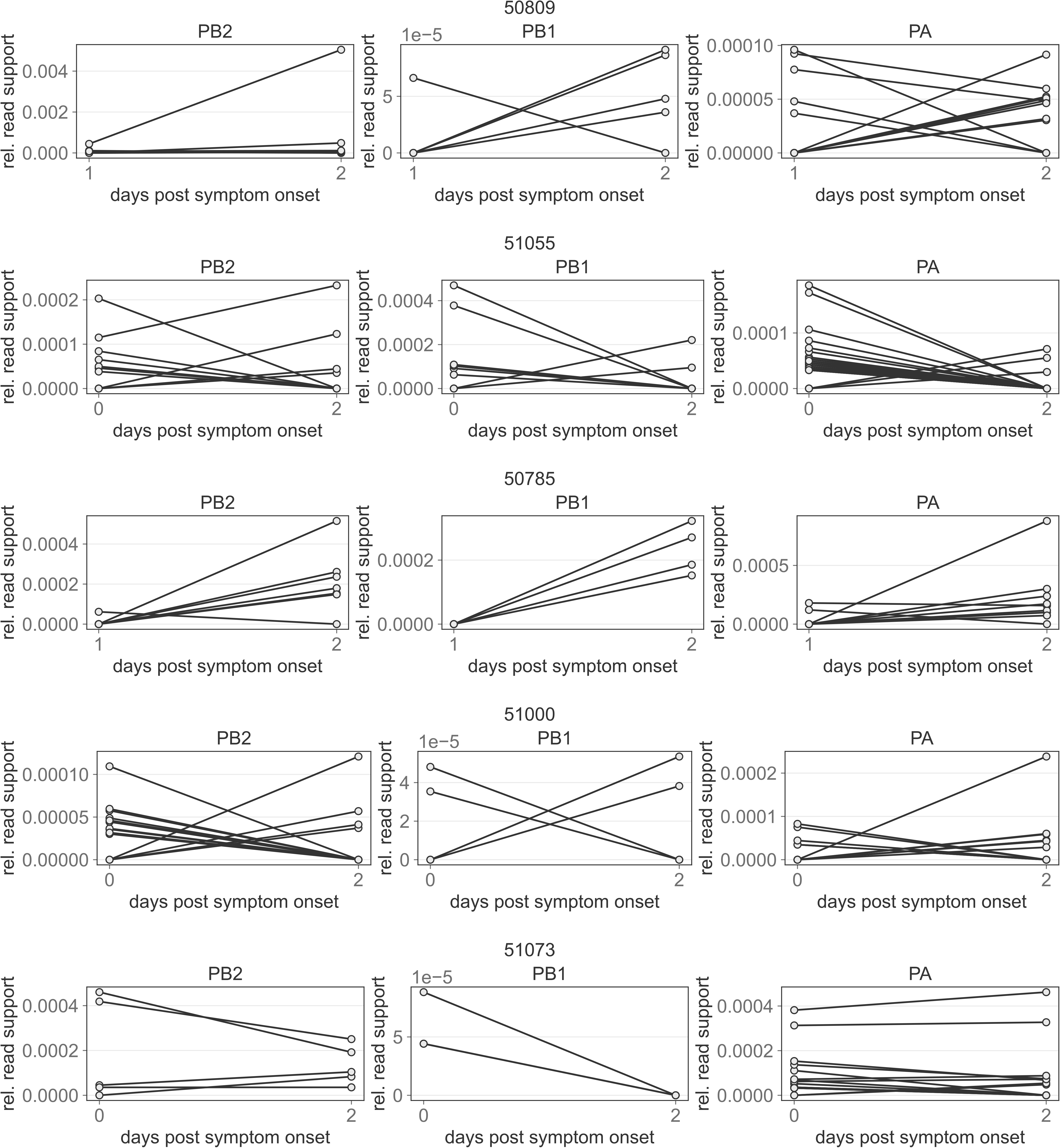
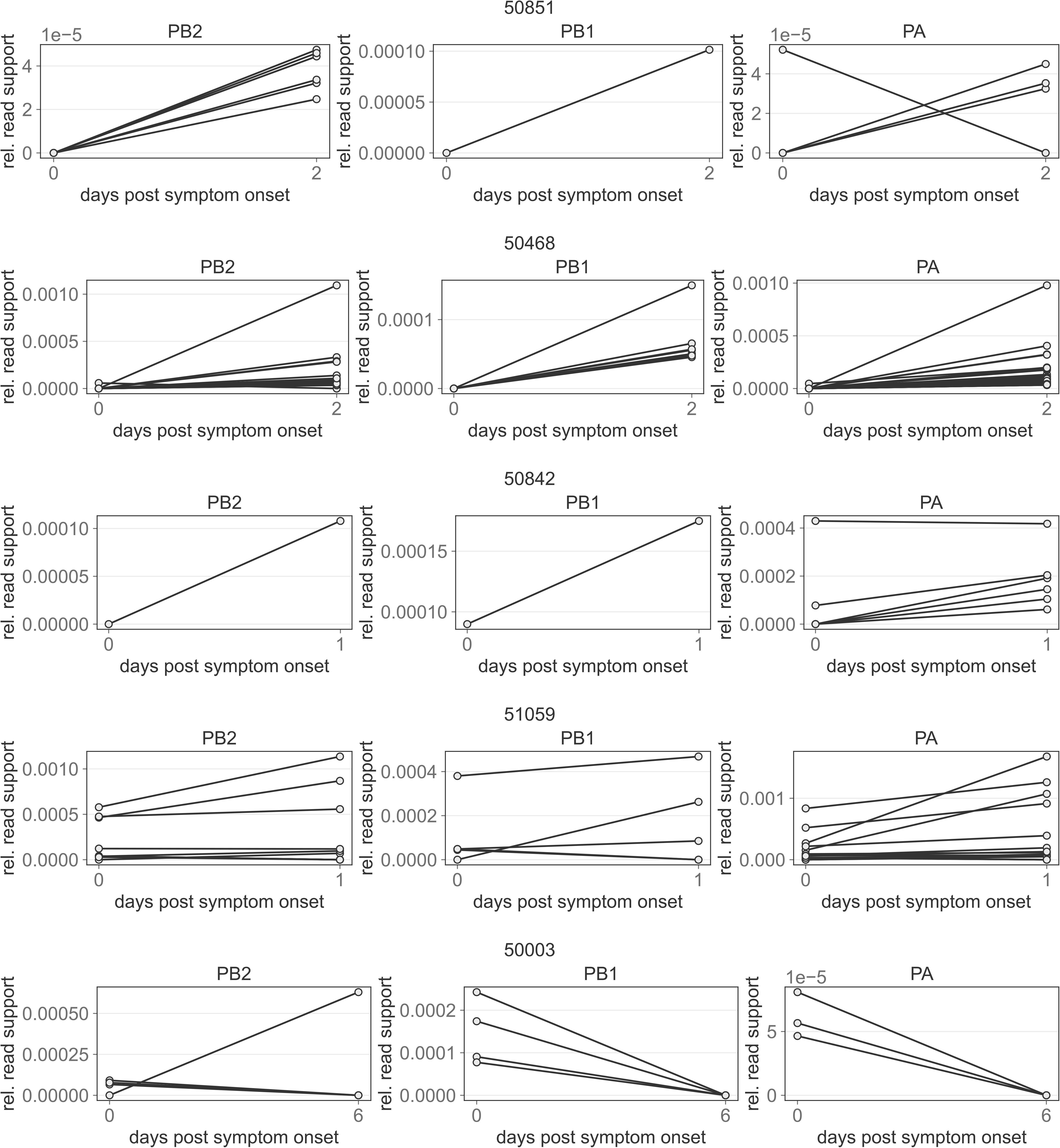
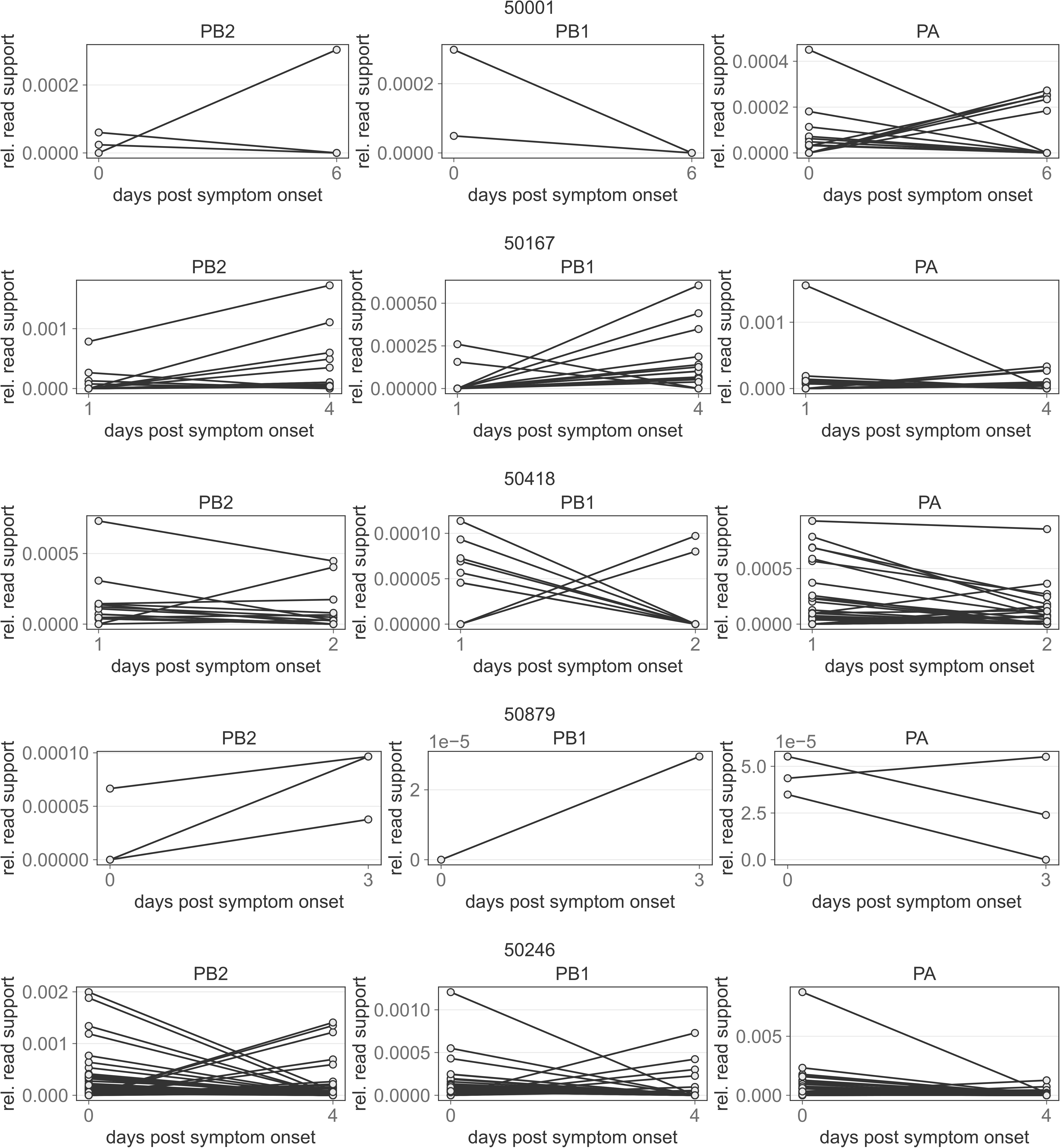
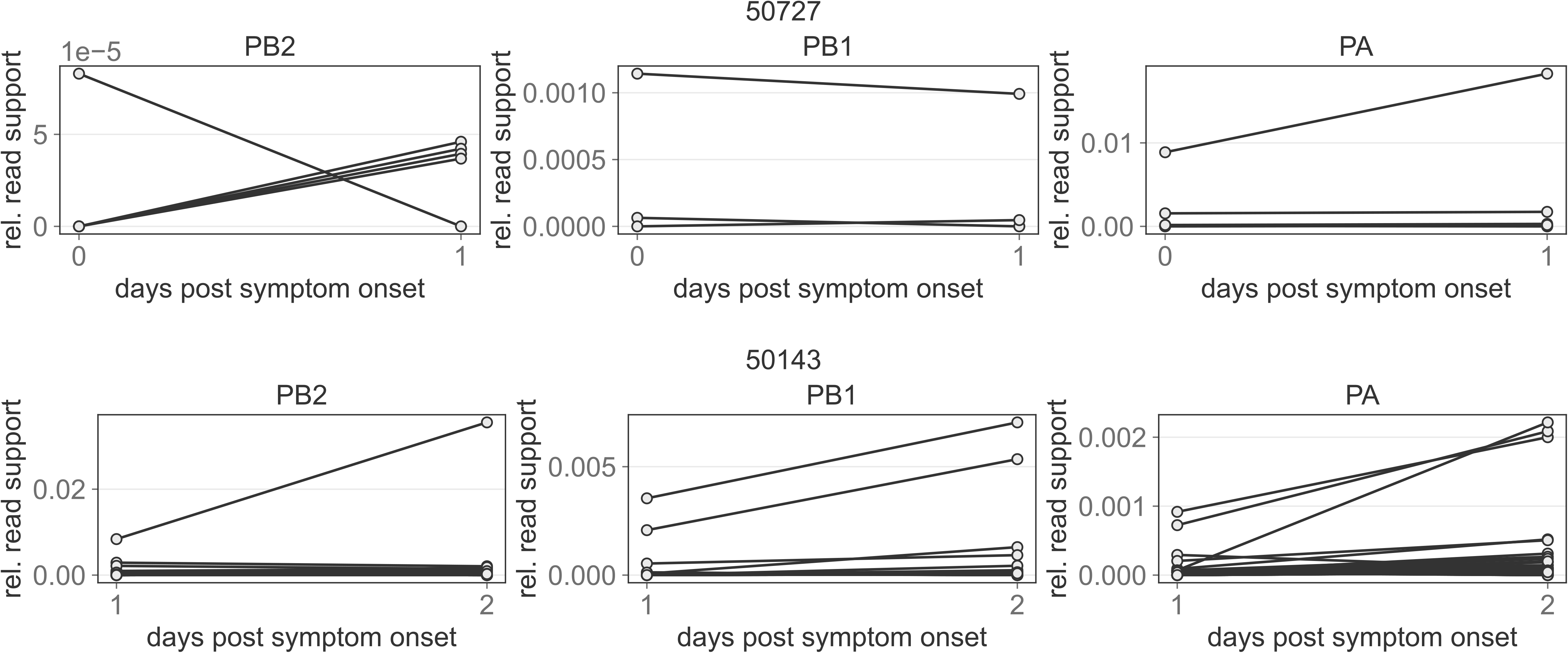
Read support of longitudinal DVGs. Relative read support of DVGs identified in the PB2, PB1, and PA segments of all individuals with longitudinal samples taken at least one day apart.

**S5 Fig.**
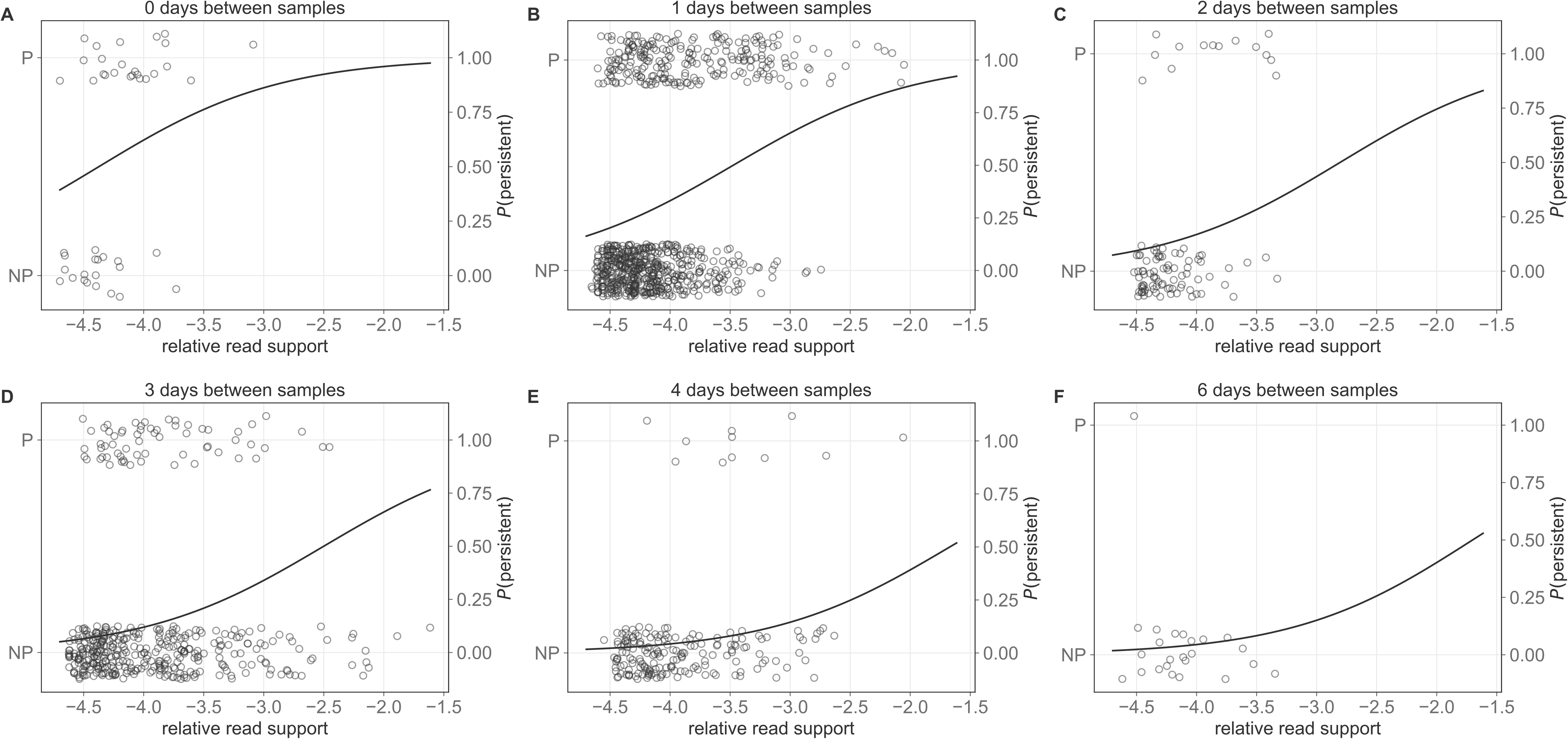
DVG persistence across time point and relative read support. Relative read support of all DVGs identified in longitudinal samples taken 0, 1, 2, 3, 4, and 6 days apart, stratified by whether those DVGs are persistent (P) or non persistent (NP) between *t*_0_ and *t*_1_. Curve is the predicted probability of persistence from a multivariate logistic regression with relative read support and time between samples (categorical) as predictors.

**S1 Table.** **DVG persistence across time points.** Logistic Regression of the probability of polymerase DVG persistence between *t*_0_ and *t*_1_ as a function of relative DVG read support.

| Variable | Coefficient | Std Err | $p$-value | 0.025 | 0.975 |
| --- | --- | --- | --- | --- | --- |
| Intercept | 7.1667 | 0.835 | 0.000 | 5.531 | 8.803 |
| $\log_{10}$ relative support | 1.9646 | 0.207 | 0.000 | 1.559 | 2.370 |

**S2 Table.** **DVG persistence across time point and relative read support.** Logistic Regression of the probability of polymerase DVG persistence between *t*_0_ and *t*_1_ as a function of time between samples and relative DVG read support.

| Variable | Coefficient | Std Err | $p$ value | 0.025 | 0.975 |
| --- | --- | --- | --- | --- | --- |
| Intercept | 5.8362 | 0.654 | 0.000 | 4.555 | 7.118 |
| $C(t_{span} = 1)$ | -1.1993 | 0.321 | 0.000 | -1.829 | -0.570 |
| $C(t_{span} = 2)$ | -2.0925 | 0.427 | 0.000 | -2.929 | -1.256 |
| $C(t_{span} = 3)$ | -2.4994 | 0.351 | 0.000 | -3.187 | -1.812 |
| $C(t_{span} = 4)$ | -3.6072 | 0.452 | 0.000 | -4.493 | -2.721 |
| $C(t_{span} = 6)$ | -3.5663 | 1.073 | 0.001 | -5.668 | -1.464 |
| $\log_{10}$ relative support | 1.3345 | 0.136 | 0.000 | 1.069 | 1.600 |

## Acknowledgments

We thank Adam Lauring, Chris Brooke, and Fadi Alnaji for helpful discussions about influenza deep sequencing and DVG identification from deep sequencing data. We thank Andrew Routh for assistance implementing the ViReMa algorithm. We thank the Koelle Lab, Daniel Weissman, Anice Lowen, Anne Piantadosi, and Tim Read for feedback on this work. Research reported in this paper was supported by the US Defense Advanced Research Projects Agency (DARPA) INTERCEPT W911NF-17-2-0034 contract (KK, MM), National Institutes of Health (NIH) National Institute of Allergy and Infectious Diseases (NIAID) Centers of Excellence for Influenza Research and Response (CEIRR) contract 93775N93021C00017 (KK), NIH NIAID F31 AI154738 (MM), and the Emory University Summer Undergraduate Research Experience (SURE) program (NB).

## Notes

https://www.ncbi.nlm.nih.gov/bioproject/?term=PRJNA412631

### Competing Interest Statement

The authors have declared no competing interest.

